# Metaplasia Enables Stomach Colonization by *Fusobacterium animalis*

**DOI:** 10.64898/2025.12.16.694801

**Authors:** C. Gómez-Garzón, Q. Chen, V.P. O’Brien, N.R. Salama

## Abstract

Infection with *Helicobacter pylori* is the major risk factor for gastric cancer worldwide; yet the exact mechanisms behind this link remain unclear. *H. pylori*-associated tissue changes often disrupt the gastric microbiome, enabling secondary gastric colonization by oral bacteria. Among these secondary colonizers, *Fusobacterium* species have documented associations with several gastrointestinal cancers. We found that both *F. animalis* and *F. nucleatum* invade cultured human gastric adenocarcinoma cells, but *F. animalis* exhibited higher adherence and invasion, and hypoxic conditions promoted higher bacterial survival. Both adherence and invasion were inhibited by exogenous GalNAc, a glycan commonly observed in membrane glycoproteins of adenocarcinoma cells, and a target of the fusobacterial adhesin Fap2. Using a mouse model of gastric metaplasia, we found that *F. animalis* colonized gastric tissue only after metaplasia onset, growing in multispecies biofilms in the mucus layer, while *F. nucleatum* colonized neither healthy nor metaplastic gastric tissue. Metaplasia led to upregulation of Gal-GalNAc in the stomach, and reduced gastric acidity allowed higher *F. animalis* loads in this model. By contrast, inflammation and the presence of *H. pylori* did not significantly influence stomach colonization by *F. animalis*. Overall, our data support a model in which *H. pylori*-induced metaplasia makes the stomach susceptible to secondary infection by another cancer-associated microbe, *F. animalis*.

## INTRODUCTION

Gastric cancer ranks as the fifth most common and lethal cancer type worldwide, with about 970,000 new cases and 660,000 new deaths in 2022, and it is estimated that the yearly incidence will reach 1.84 million new cases by 2050 (1). Current estimates suggest that 77 % of gastric cancer cases are attributable to infection with the bacterium *Helicobacter pylori*, making it the major risk factor for this malignancy (2). Although the exact mechanism by which *H. pylori* infection causes cancer development remains unclear, an *H. pylori*-led cascade of cancer associated tissue changes has been extensively documented (3). Initially, *H. pylori* triggers gastritis, chronic inflammation of the gastric epithelium, which may progress to atrophic gastritis, defined by the loss of the acid- and digestive enzyme-producing parietal and chief cells, respectively. Atrophic gastritis may further progress to metaplasia and dysplasia, and ultimately, cancer onset in 1 – 3 % of *H. pylori* infected individuals (4–6).

High acidity and *H. pylori* infection shape the gastric microbiome (7). In individuals infected with *H. pylori*, about 50 % of the world population (6), this bacterium is the most prevalent bacterial species in the stomach. However, increased pH associated with atrophic gastritis leads to gastric dysbiosis, characterized by a significant reduction in *H. pylori* abundance and increased colonization by species from the oral microbiome, including genera such as *Fusobacterium*, *Prevotella*, and *Streptococcus* (8–11). Thus, bacteria other than *H. pylori* may play a role in gastric cancer development.

Among these genera, *Fusobacterium* species have been reported as tumor-associated bacteria in cases of breast (12, 13), pancreatic (14, 15), colorectal (16–18), nasopharyngeal (19), oesophagogastric (20), and gastric cancer (8, 11, 21–23). *F. nucleatum* is the most studied species of this genus given its prevalence in the oral cavity and its association with oral pathologies. However, it has been reported that different species possess different virulence factors and may be distinctly associated with different pathologies (24). Regarding cancer, *F. animalis* (also referred to as *F. nucleatum* subsp. *animalis* C2 in the literature) is the most commonly found *Fusobacterium* species in colorectal cancer cases (16), which remains the most studied case of *Fusobacterium*-associated malignancy. In colorectal cancer, Fusobacteria exhibit immunosuppressive and proinflammatory activities, promote metastasis (25, 26) and migrate with metastatic cells to the liver (27). In gastric cancer, evidence is scarcer, albeit Fusobacteria have documented associations with poorer prognosis (23, 28–30) and may promote gastric tumor progression (28).

In the current study, we assessed the relative capabilities of *F. animalis* and *F. nucleatum* to infect gastric cancer cells, using the AGS cell line. To explore cellular and tissue features that promote gastric tissue colonization, we used an inducible transgenic mouse model of gastric metaplasia. Our results show that *F. animalis* is better adapted than *F. nucleatum* to colonize both gastric cancer cells and metaplastic stomach tissue, with the onset of metaplasia being a critical factor enabling stomach colonization.

## RESULTS

### *F. animalis* invades human adenocarcinoma cancer cells *in vitro* under hypoxic conditions

Fusobacteria are facultative intracellular pathogens that have been reported as members of the intratumoral microbiota in several types of cancer, including gastric cancer (8, 11, 20, 22, 23). To test whether *Fusobacterium* species can invade gastric cancer cells (**Fig. 1A**), we performed co-culture with AGS cells, a commonly used gastric adenocarcinoma cell line (31). Given recent work suggesting distinct species colonize the oral cavity, but *Fusobacterium animalis* is prevalent in colorectal cancer tumors (16), we first tested the *F. animalis* strain SB010, originally isolated from a colorectal cancer case (27). We performed co-culture under microaerobic conditions (10 % O_2_, 10 % CO_2_) or hypoxia (1.0 % O_2_, 10 % CO_2_), considering the anaerobic lifestyle of this bacterium. In short, we inoculated AGS monolayers with *Fusobacterium* at a multiplicity of infection (MOI) of 0.1 and incubated the co-culture under microaerobiosis or hypoxia for 4 h and 24 h. After incubation, we removed unbound and loosely bound bacteria by repeated washes and quantified cell-associated bacteria (*i.e.,* adhered and intercellular). Separately, we also quantified intracellular bacteria by killing all extracellular bacteria with metronidazole, an antibiotic that is not membrane-permeable and hence does not affect internalized bacteria. As shown in **Fig. 1B**, we found that hypoxic conditions allowed for more robust adherence and invasion at 4 h, with an approximately 6-fold difference in colony forming units (CFU) compared to the microaerobic condition for both cell-associated (*p* = 0.033) and intracellular bacteria (*p* = 0.048). On average, 19 % and 25% of the initial *F. animalis* inoculum remained viable after the 4 h of co-culture under microaerobic and hypoxic conditions, respectively (that is, both host-associated and free bacteria). Out of these viable bacteria, the ratio of host-associated bacteria was 15 % and 69 % for the same conditions. Notably, after 24 h under microaerobic conditions, we did not recover detectable viable *F. animalis* CFUs adhered or intracellular. Only hypoxia enabled *F. animalis* survival after 24 h of co-culture with AGS cells, with 3.0 % of the initial inoculum remaining viable, and 70 % of these viable bacteria corresponding to host-associated bacteria.

**Figure 1.**
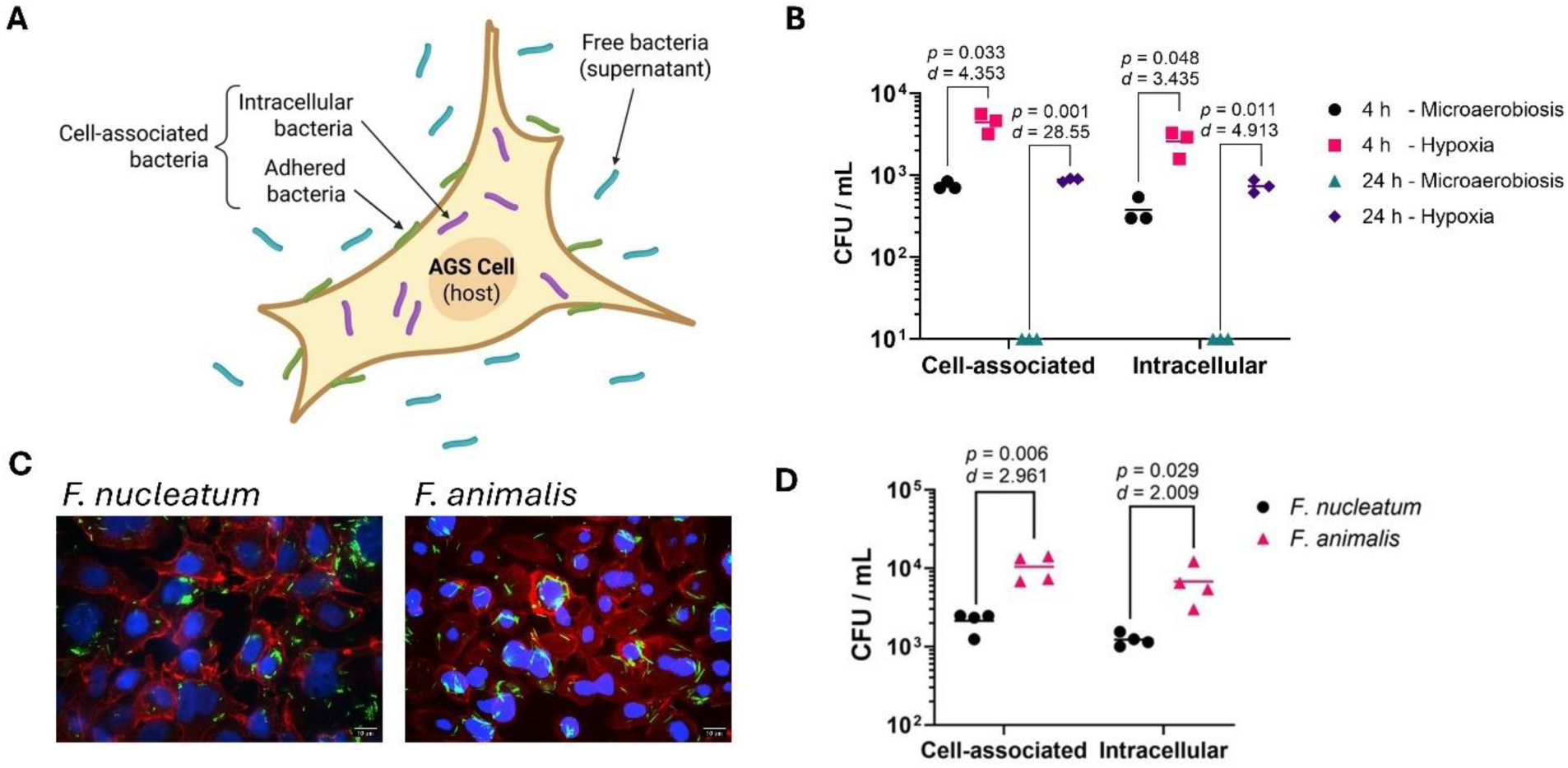
Hypoxic conditions facilitate *F. animalis* adherence to and invasion of AGS cells. **A)** Schematic representation of the bacterial subpopulations quantified during co-culture with AGS cells. Free bacteria were washed away, and adhered bacteria were killed with metronidazole to quantify intracellular bacteria. CFU counts from replicate wells without metronidazole were plated to enumerate total cell-associated bacteria. Created with BioRender.com **B)** *F. animalis* survival at 4 h or 24 h during co-culture with AGS cells under microaerobic (10 % O_2_, 10 % CO_2_) or hypoxic (1.0 % O_2_, 10 % CO_2_) conditions, Multiplicity of infection (MOI) of 0.1. **C)** Fluorescence microscopy of AGS cells infected with *F. nucleatum* (left) or *F. animalis* (right). Nuclei, in blue (DAPI); F-actin, in red (Phalloidin-Alexa Fluor 555); and *Fusobacterium*, in green (CFSE). MOI = 10. **D)** *F. nucleatum* and *F. animalis* co-culture with AGS cells under hypoxia for 4 h with MOI = 0.1. Panels B and D show data from a representative experiment out of three independent replicates. Horizontal lines represent the mean of each group, *p-*values correspond to unpaired, two-tailed t-tests between groups, and *d*-values represent Cohen’s d-value for the effect size. Zeroes were plotted at the limit of detection (10 CFU / mL).

We next compared *F. animalis* SB010 to the type strain *F. nucleatum* ATCC 23726, a representative oral fusobacterial strain, for adherence and intracellular invasion under hypoxia after 4 h. Although both species were able to invade AGS cells, as seen by fluorescence microscopy (**Fig. 1C**), we found that *F. animalis* exhibited higher adherence and invasiveness, with 5-fold higher CFUs for both host-associated (*p* = 0.006) and intracellular bacteria (*p =* 0.029) (**Fig. 1D**). Overall survival of *F. animalis* and *F. nucleatum* under these conditions was estimated at 24 % and 4.4 % of the initial inoculum, respectively. Taken together, these results indicate that conditions of reduced oxygen pressure facilitate the study of intracellular invasion by *F. animalis*, and that this species exhibits higher adherence, invasion, and survival capabilities in AGS cells than *F. nucleatum*.

### *F. animalis* intracellular invasion of AGS cells is inhibited by GalNAc

An increasing body of evidence suggests that D-galactose-β(1–3)-N-acetyl-D-galactosamine (Gal-GalNAc) is a target on host cell surface that Fusobacteria utilize to invade cancer cells via recognition by the bacterial adhesin Fap2 (32). To assess the importance of Gal-GalNAc binding in infection of gastric cancer cells, we incubated *F. animalis* with 25 mM GalNAc for 30 min prior to infection, and observed significantly reduced *F. animalis* adherence (*p* = 0.001) and invasion (*p* = 0.001) of AGS cells compared to the control incubated with phosphate saline buffer (PBS) (**Fig. 2A**). We verified that GalNAc does not affect the viability of *F. animalis* (**Supplementary Fig. 1**). Interestingly, we also observed that there were more viable bacterial cells in the supernatants of the GalNAc-treated group than in the control group (**Fig. 2B**), which further confirms that exogenous GalNAc prevents *F. animalis* from binding the AGS cells without affecting their viability. AGS cells produce Gal-GalNAc as measured by binding of FITC-conjugated peanut agglutinin (PNA), a Gal-GalNAc-binding lectin (**Fig. 2C**).

**Figure 2.**
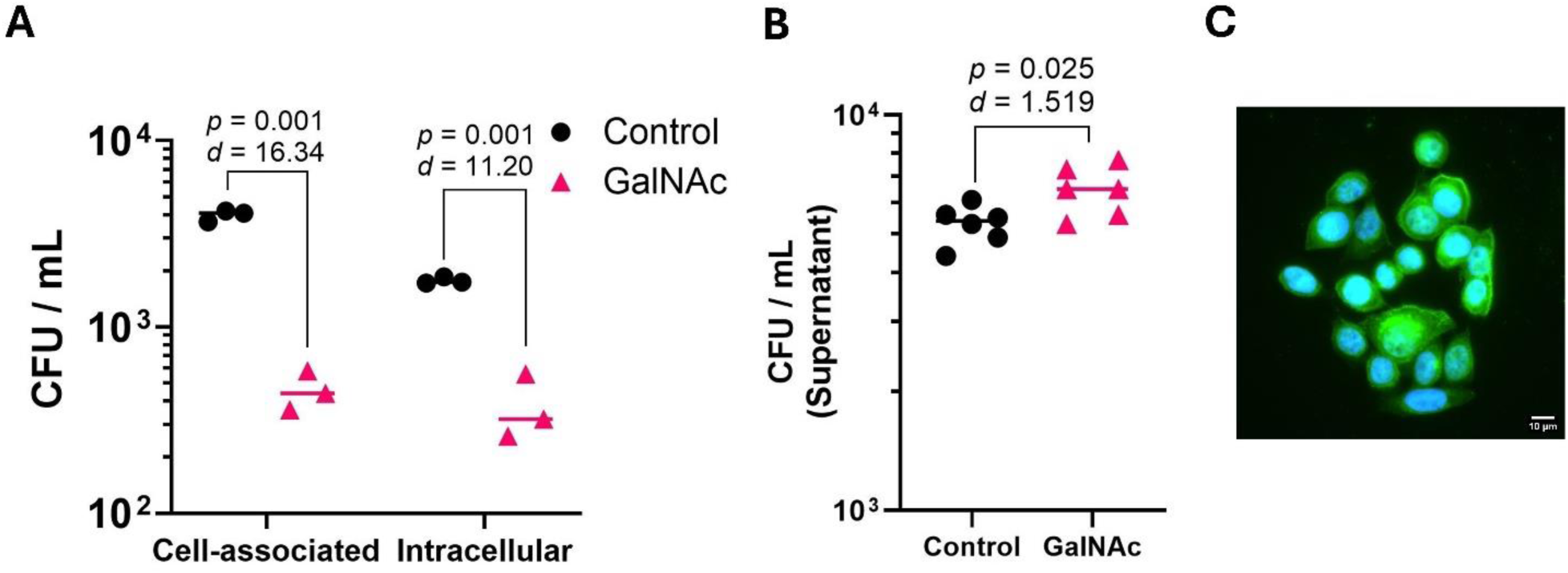
*F. animalis* adherence and invasion of AGS cells are blocked by GalNAc. **A)** *F. animalis* cells were pretreated with GalNAc (25 mM) or PBS (control) for 30 min. Metronidazole (300 µM, 90 min) was used to differentially quantify intracellular bacteria. Titers of the indicated cell fraction after 4 h of co-culture are shown. **B)** *F. animalis* titers in the supernatants of the co-cultures with AGS cells shown in panel A. **C)** AGS cells stained with FITC-conjugated PNA (binds Gal-GalNAc) in green. Nuclei are shown in blue (DAPI). For panels A and B, plotted data show a representative experiment out of three independent replicates. Horizontal lines represent the mean of each group. The *p-*values correspond to unpaired, two-tailed t-tests between groups, and *d*-values represent Cohen’s d estimate for effect size.

### Gastric metaplasia onset promotes *F. animalis* infection in a mouse model

Motivated by the widely documented association of Fusobacteria with cancer cells (33–35), together with our results from the *in vitro* AGS cells model, we wondered whether Fusobacteria can also infect the gastric environment during precancerous stages. To address this question, we leveraged a mouse model of gastric intestinal metaplasia (GIM), *Mist1-CreERT2^Tg/+^ LSL-Kras*^G12D^*^Tg/+^* (herein referred to as *Mist1-Kras* mice) (36) that our lab has employed to study the effect of *H. pylori* chronic infection on gastric preneoplastic progression (37, 38). These mice harbor an allele encoding Kras^G12D^ (a constitutively active variant of Kras) that can be induced through excision of a stop codon flanked by *loxP* sites via tamoxifen allosteric activation of a CreERT2 enzyme under control of the *Mist1* promoter. In the stomach, Mist1 is expressed in gastric chief cells and a subset of gastric progenitor cells. When induced with tamoxifen (*Kras*+), these mice develop metaplasia with temporally varying features, including spasmolytic polypeptide-expressing metaplasia (SPEM) within four weeks, which displays intestinal characteristics (SPEM-IC) by six to eight weeks, and finally intestinal metaplasia (IM) by the 12^th^ week (**Fig. 3A**).

**Figure 3.**
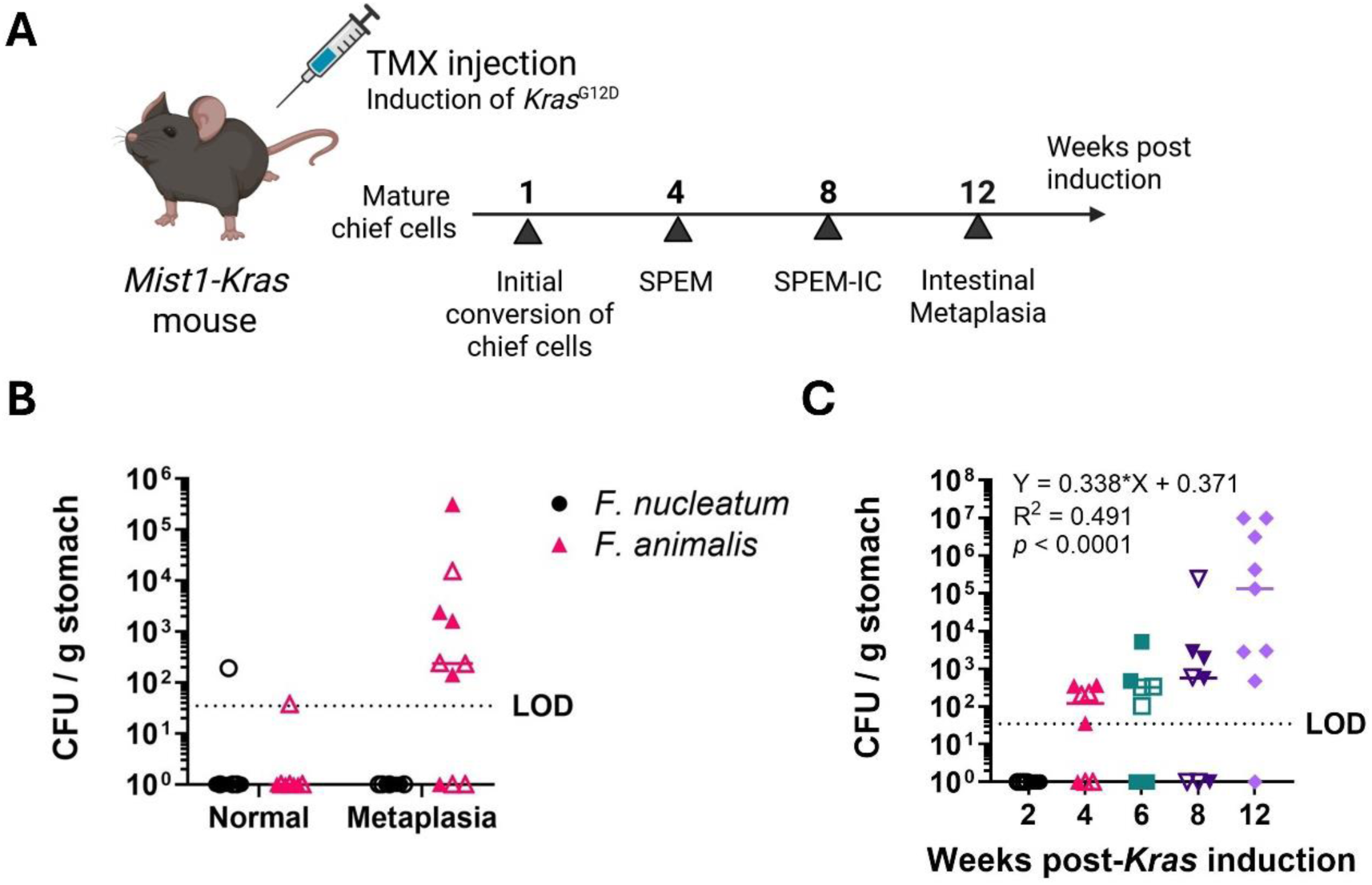
Metaplasia onset promotes *F. animalis* gastric infection. **A)** Constitutive expression of *Kras*^G12D^ in chief cells was induced in *Mist1-Kras* mice via tamoxifen injection, leading to the depicted preneoplastic cascade. SPEM = Spasmolytic Peptide Expressing Metaplasia, IC = intestinalizing characteristics. Panel adapted from Choi *et al.* 2016 (36) using BioRender.com **B)** Gastric *F. animalis* titers recovered one week post-infection from mice at 6 weeks post-tamoxifen (Metaplasia) vs. sham-induced mice (Normal). **C)** Gastric *F. animalis* loads recovered from mice after the indicated time post-*Kras* induction. Ordinary one-way ANOVA on log-transformed data, *p*-value < 0.0001, F = 8.627. The equation, R^2^, and *p*-value (testing slope equal to zero by F-test) shown in the plot correspond to the log-linear regression for CFU vs. weeks post-*Kras* induction. Data plotted in panels B and C were obtained from two independent experiments, indicated with open and filled symbols, except for the 12-week timepoint in panel C. Horizontal lines represent the median of each group.

We orally infected mice 6 weeks post-*Kras* induction (after SPEM onset, hereafter referred to as metaplasia) and sham-induced mice (control) with either *F. animalis* or *F. nucleatum*. Gastric fusobacterial loads were quantified one week post-infection. Except for a single outlier, *F. nucleatum* was unable to colonize the mouse stomach, regardless of the metaplastic status. By contrast, *F. animalis* robustly colonized the metaplastic stomach (7/10 animals infected with a median of 2.37 × 10^2^ CFU / g stomach) but not the normal stomach (sham-induced, 1/8 mice infected, *p* = 0.0395) as shown in **Fig. 3B**.

To assess stomach colonization by *F. animalis* throughout the preneoplastic cascade, we infected *Kras*+ mice with *F. animalis* at different time points after *Kras* induction. Mice were orally infected with *F. animalis* at 2, 4, 6, 8, and 12 weeks post-induction, corresponding to the progression from initial chief and progenitor cell conversion to the onset of intestinal metaplasia (**Fig. 3A**), and gastric *F. animalis* colonization was quantified at 1 week post-infection. We did not detect infection 2 weeks after *Kras* induction. Starting at 4 weeks after *Kras* induction, we recovered *F. animalis,* and loads increased at each timepoint thereafter following a log-linear relationship (**Fig. 3C**). Ordinary one-way ANOVA on log-transformed data had a *p*-value < 0.0001 and F = 8.63, and post-hoc Tukey’s tests for multiple comparisons showed significant differences only for the 12-week group compared to all the others (*p* < 0.0001, and *p =* 0.003, 0.025, and 0.041 vs the 2, 4, 6, and 8-week groups, respectively). Thus, in mice, *F. animalis* can colonize the metaplastic, but not the healthy, stomach, and the *F. nucleatum* type strain cannot colonize either normal or metaplastic gastric tissue.

### Elevated pH facilitates *F. animalis* gastric colonization

We next explored which features of the metaplastic environment promote colonization by *F. animalis*. Gastric metaplasia is characterized by increased stomach pH (due to loss of parietal cells), inflammation, and altered gene expression, mucin production, and cell surface glycosylation (39, 40). We evaluated whether increased pH alone (*i.e.*, without metaplastic alterations) would allow *F. animalis* to colonize the gastric niche. We administered omeprazole, a widely used proton pump inhibitor, daily to WT mice starting three days before infection with *F. animalis*. The infection was allowed to progress for one week, during which we continued the daily omeprazole administration to sustain a less acidic gastric environment in these mice. At the end of this experiment, we did not detect any instances of *F. animalis* gastric infection in either the treatment or the control group, which received only the drug vehicle (40 % v/v PEG 400) (**Fig. 4A**).

**Figure 4.**
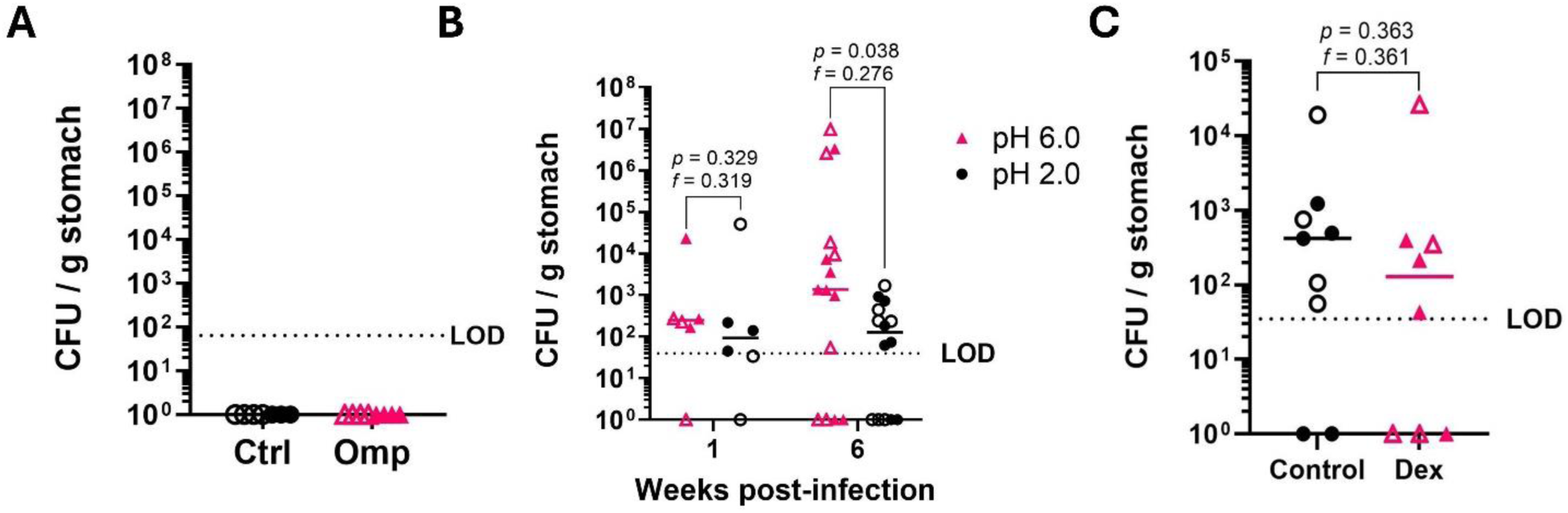
Impact of gastric pH and inflammation on *F. animalis* colonization of normal and metaplastic stomach tissues. **A)** Gastric loads of *F. animalis* 1 week post-infection in WT mice treated with omeprazole (Omp) or the drug vehicle (40 % PEG 400) as control. **B)** Gastric loads of *F. animalis* 1 and 6 weeks post-infection recovered from *Mist1-Kras* mice 6 weeks post-tamoxifen induction receiving either neutral (pH 6) or acidified (pH 2) drinking water. **C)** Gastric loads of *F. animalis* in *Mist1-Kras* mice 6 weeks after tamoxifen induction with or without dexamethasone sodium phosphate (Dex) added to the drinking water to suppress inflammation*. p*-values correspond to non-paired Mann-Whitney U tests; and the *f*-values, to the common language effect size. Data were obtained from two independent replicates of each experiment, indicated with open and filled symbols, using 3 – 7 mice per group. Horizontal lines represent the median values, and LOD stands for limit of detection.

Additionally, we tested whether elevated pH facilitates *F. animalis* infection of GIM stomach tissue. We compared gastric *F. animalis* titers from *Mist1-Kras* mice infected 6 weeks after tamoxifen administration, given either acidified drinking water (pH ≈ 2) or non-acidified autoclaved drinking water (pH ≈ 6), which is the standard condition used in all other experiments reported herein. At one week post-infection, the group receiving acidified drinking water exhibited a modest decrease in *F. animalis* loads (**Fig. 4B**). This difference became more pronounced at 6 weeks post-infection, when the acidified drinking water group exhibited significantly lower gastric *F. animalis* loads (**Fig. 4B**). Collectively, these results indicate that a more neutral gastric environment facilitates *F. animalis* infection of metaplastic stomach tissue but does not allow colonization of normal stomach tissue.

Given the strong association of Fusobacteria with inflamed tissues, we also tested whether the metaplasia-driven inflammation might promote fusobacterial infection of the stomach using the immunosuppressive glucocorticoid dexamethasone. Mist1-Kras mice were immunosuppressed with oral dexamethasone sodium phosphate *ad libitum* in the drinking water, starting 3 weeks after tamoxifen induction. At 6 weeks after induction, mice were orally gavaged with *F. animalis*, and infection was assessed after one week, while dexamethasone treatment was continued. Oral dexamethasone produces immune attenuation in mice (38), and animalis in these experiments exhibited the classical weight loss and reduced spleen mass induced by this drug (Supplementary Fig. 2). The distribution of *F. animalis* loads recovered from these mice was not different from the control or that observed in the previous experiments, indicating that the immunosuppression induced by the dexamethasone did not affect *F. animalis* infection of GIM stomach tissue (**Fig. 3C**).

### Metaplasia onset leads to increased gastric Gal-GalNAc levels

Since neither inflammation nor pH appeared to be a major factor driving *F. animalis* colonization of metaplastic gastric tissue, we considered expression of known *F. animalis* cellular receptors. Given our results that Gal-GalNAc promotes *F. animalis* invasion of a gastric cancer cell line (**Fig. 3**), we tested whether the onset of GIM in *Mist1-Kras* mice leads to increased production of Gal-GalNAc. We used FITC-labeled PNA to survey the presence of Gal-GalNAc in gastric tissues of mice at different points of the preneoplastic progression (**Fig. 5**). The stomach of the control group showed modest production of Gal-GalNAc, mainly limited to the region comprising the transition between the antrum and the corpus and confined to the very tops and base of the glands. By contrast, we observed higher Gal-GalNAc in the corpus of the induced *Mist1-Kras* mice on cells comprising the top1/3-1/2 of the glands. Gal-GalNAc upregulation was observed as early as 2 weeks after *Kras*-induction and sustained through week 12. These data suggest that disruption of protein glycosylation, leading to Gal-GalNAc upregulation, accompanies initial chief cell conversion (2^nd^ week) during the Kras-driven preneoplastic cascade, and continues until the intestinal metaplasia stage. Notably, Gal-GalNAc upregulation preceded susceptibility to *F. animalis* infection, which we only detected 4 weeks post-induction (**Fig. 3C**). Thus, while Gal-GalNAc may promote *F. animalis* invasion in cancer cells, additional features of the metaplastic stomach must contribute to gastric tissue surface colonization.

**Figure 5.**
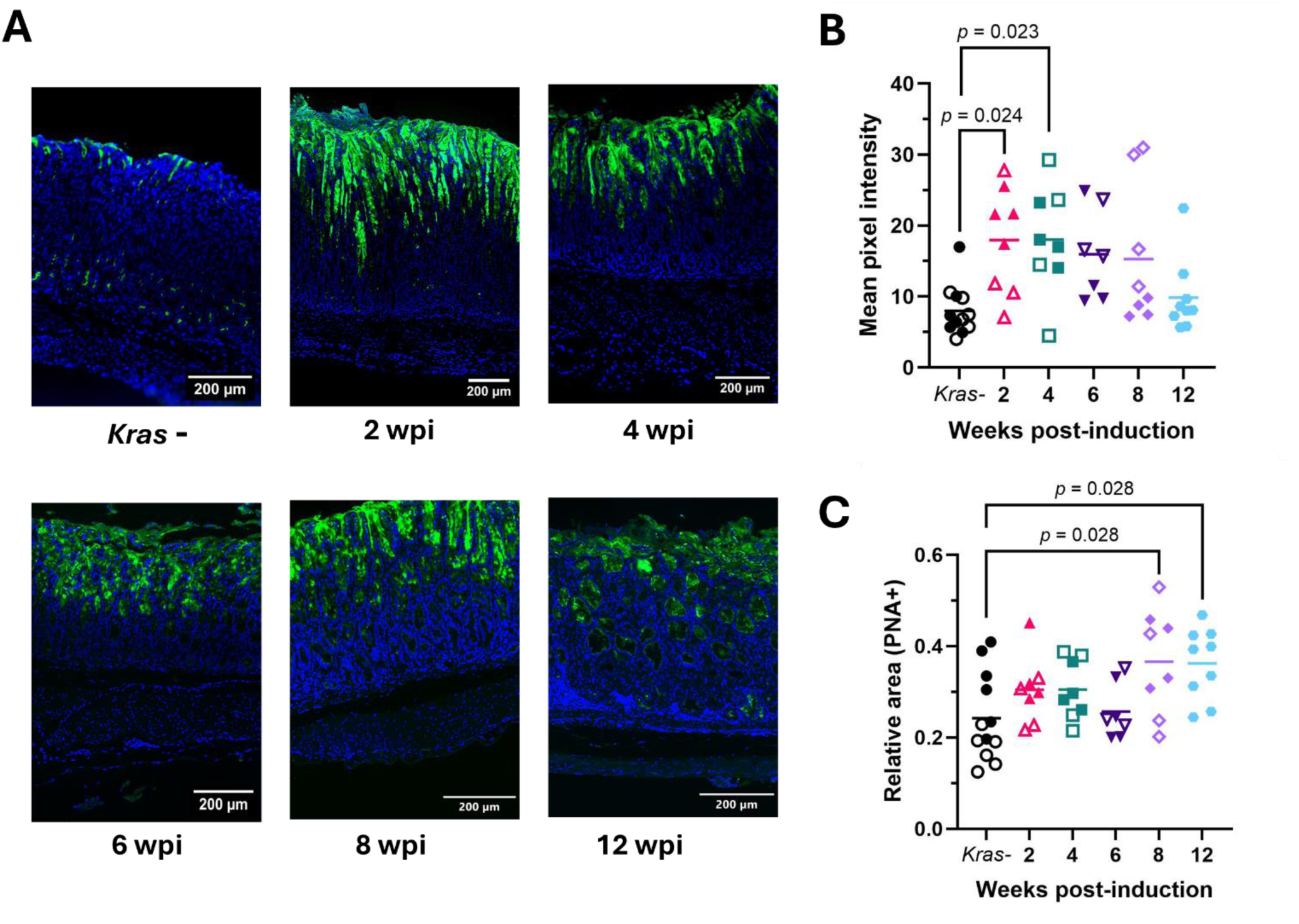
Gal-GalNAc production in mouse stomach tissues increases during *Kras*-driven preneoplastic progression. **A)** Gal-GalNAc visualized with FITC-conjugated peanut agglutinin (PNA, green) and nuclei stained with DAPI (blue). The images correspond to representative sections of stomach corpora of mice at different points after *Kras* induction (wpi = weeks post-induction). Uninduced (*Kras*-) mice were included as controls. **B)** Quantification of the mean FITC pixel intensity and **C)** FITC relative area, relative to DAPI staining, in mouse gastric corpora vs. weeks post-*K*ras induction. Data were obtained from two independent experiments, indicated with open and filled symbols, except for the 12-week timepoint. Horizonal lines show mean values. Statistical analysis was conducted by an ordinary one-way ANOVA (B: *p* = 0.005, F = 3.893 and C: *p* = 0.0105, F = 3.406), and the shown *p*-values correspond to significant (< 0.05) post-hoc comparisons with a Tukey’s test.

In addition, we surveyed Gal-GalNAc production in a previously described tissue microarray (TMA) comprising 2-mm tissue cores from a cohort of 47 gastric cancer patients from medical centers in the U.S. Pacific Northwest (**Figure 6**) (38). Tissue cores from each patient include regions of superficial cancer, deep cancer, and non-neoplastic epithelium adjacent to cancerous lesions. To test whether Gal-GalNAc display correlates with fusobacterial infection, we used droplet digital PCR (ddPCR) to detect Fusobacteria (including *F. animalis* and *F. nucleatum*) in these tissues. However, we did not find a significant correlation between Fusobacteria status and Gal-GalNAc production, indicating that Gal-GalNAc may not be the only determining factor for stomach colonization by Fusobacteria in either metaplastic or gastric cancer tissue.

**Figure 6.**
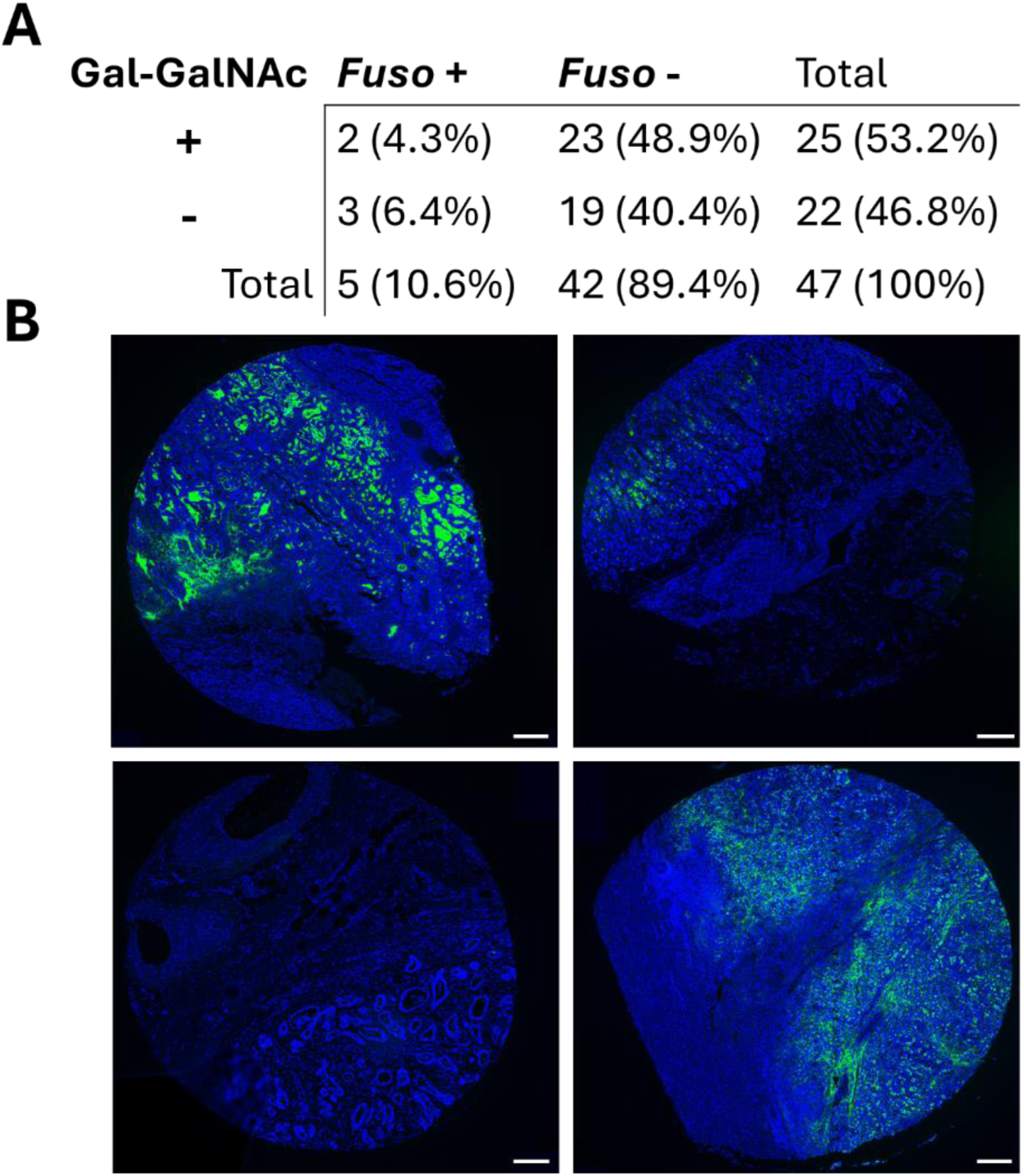
Gal-GalNAc and Fusobacteria status in a gastric cancer TMA. **A)** Contingency table showing the number of cases, and percentage of total cases, classified according to Gal-GalNAc detection (+/− using FITC-PNA staining) and fusobacterial infection (*Fuso* +/−, detected by ddPCR). Data were analyzed with a two-sided Fisher’s exact test, *p* = 0.654. **B)** Representative images from TMA cores. From left to right and top to bottom: superficial cancer (Fuso+, Gal-GalNAc+), non-neoplastic epithelium adjacent to cancer (Fuso-, Gal-GalNAc+), deep cancer (Fuso-, Gal-GalNAc-), and deep cancer (Fuso-, Gal-GalNAc+). DAPI (nuclei) is shown in blue and FITC-PNA (Gal-GalNAc), in green. Scale bars = 200 µm.

### The presence of *H. pylori* neither prevents nor promotes *F. animalis* colonization of GIM stomach tissues

*H. pylori*-driven chronic inflammation leads to gastric microecological dysbiosis, which starts during precancerous stages and exhibits a distinct profile in the cancer stage (9, 11, 41, 42). *H. pylori*-associated gastric dysbiosis is often characterized by an increase in microbial diversity in the stomach, including elevated abundance of oral species such as Fusobacteria, and reduced prevalence of *H. pylori* (11, 22, 43–45). To gain insight into possible interactions between *H. pylori* and *F. animalis*, we evaluated whether these species can co-infect the stomach of *Mist1-Kras* mice. First, we inoculated *Mist1-Kras* mice with *H. pylori* concomitantly with tamoxifen induction. After 6 weeks, we infected these mice (*H. pylori*-infected, *Kras*+) with *F. animalis*, and quantified gastric CFUs of both species one week later. Our results indicate that *F. animalis* can establish an infection in GIM stomach tissues even if *H. pylori* is present (**Fig. 7A**).

**Figure 7.**
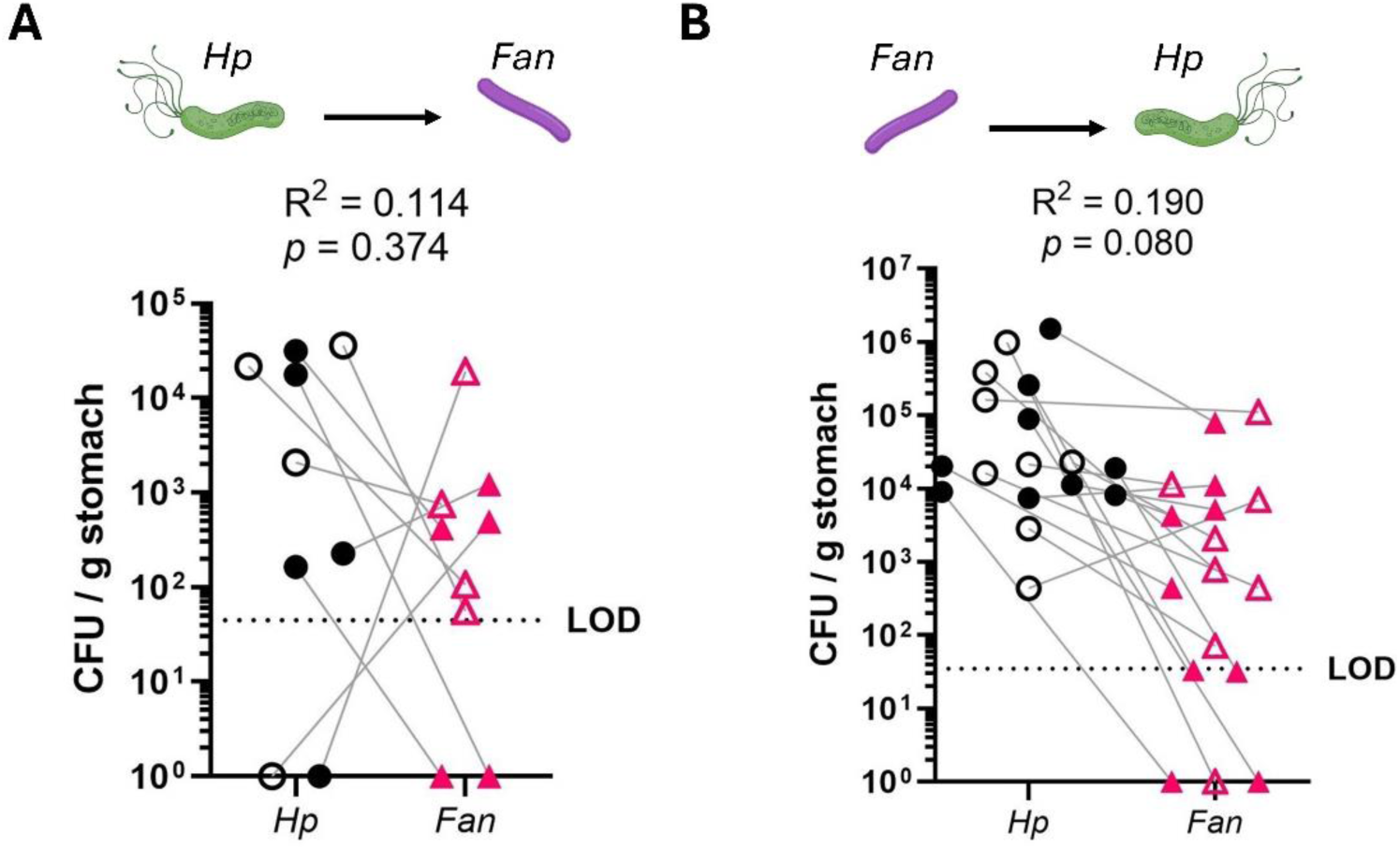
*H. pylori* (*Hp*) and *F. animalis* (*Fan*) co-infection assays in the *K*ras+ mouse model. **A)** *H. pylori*-infected, *Kras+* mice were inoculated with *F. animalis*, and bacterial loads were quantified one week later. **B)** *F. animalis*-infected, *Kras+* mice were inoculated with *H. pylori,* and bacterial loads were quantified 6 weeks post-infection. The plotted data were obtained from two independent experiments, indicated with open and filled symbols, using 4 - 9 mice per group, and shown with open and filled symbols. LOD stands for limit of detection. R^2^ corresponds to the coefficient of determination of linear correlation analyses for *Fan* vs *Hp*, and *p-*values correspond to F tests for the null hypothesis of the slope equaling zero. Experimental schematic created with BioRender.com.

Next, we tested whether an established *F. animalis* infection prevents *H. pylori* from colonizing and persisting in the metaplastic stomach, which could additionally contribute to the concomitant decrease of *H. pylori* during gastric dysbiosis. We induced *Mist1-Kras* mice with tamoxifen and orally infected them with *F. animalis* at the 6^th^ week post-induction. One week after infection, we inoculated these mice (*F. animalis*-infected, *Kras*+) with *H. pylori* and allowed the co-infection to progress for 6 weeks. After this period, the gastric bacterial loads for each species were determined. Similar to the experiment above, our results indicate that both bacterial species could establish a co-infection lasting six weeks (**Fig. 7B**), suggesting that co-infections are possible and stable. In neither order of addition did bacterial titers show a significant correlation between species.

In summary, our co-infection assays in the mouse model revealed no evidence of an inhibitory interaction between *H. pylori* and *F. animalis* in colonizing GIM stomach tissues.

### *F. animalis* colonizes the mucus layer of the metaplastic mouse stomach and triggers the production of proinflammatory and chemotactic cytokines

To study the interaction between the host and *F. animalis* in the context of gastric preneoplastic progression, we utilized fluorescent *in situ* hybridization (FISH) to localize *F. animalis* in the gastric tissue of *Kras*+ mice via confocal microscopy. We did not observe signs of intracellular *F. animalis*; instead, we found that *F. animalis* localized in the corpus mucus layer. Notably, *F. animalis* signal colocalized with that from a 16S probe for Eubacteria, suggesting that *F. animalis* is colonizing the gastric mucus in association with other bacteria (**Fig. 8 and Supplementary Figs. 3 and 4**).

**Figure 8.**
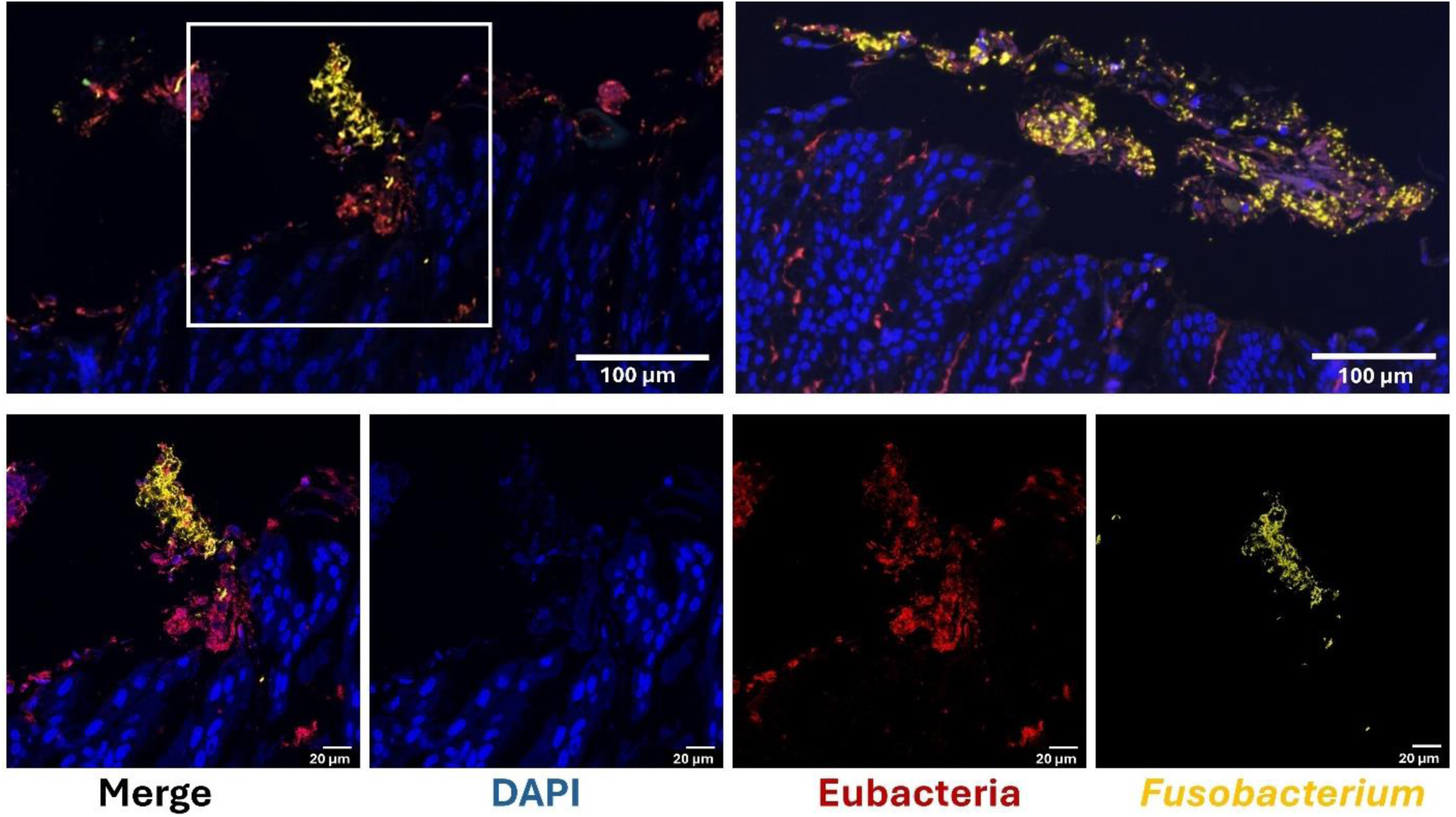
*F. animalis* proliferates in discrete foci within the gastric mucus layer of *Kras*+ mice. The top panels show representative foci located in corpus sections imaged by fluorescence *in situ* hybridization (FISH). The bottom panels present the individual separated channels of the corpus region outlined by the white box: DAPI (DNA, in blue), eubacterial 16S rRNA probe (red), and *Fusobacterium*-specific 23S rRNA probe (yellow).

Having established the contact between *F. animalis* and its host as located on the gastric epithelial surface, we sought to assess how the host responds to the infection. We surveyed the concentration of 32 different proinflammatory and chemotactic cytokines (representing the major families of these signaling molecules) in stomach homogenates of *Kras*+ mice infected with *F. animalis* for 6 weeks. As a control, the same measurements were made in homogenates from mock-infected animals. Measurements for IL-13 fell below the standard curve range. For 15 out of the remaining 31 quantified cytokines, the infected group exhibited a bimodal distribution, in which the average cytokine concentration of one subpopulation was significantly higher than that of the mock-infected group and/or the other subpopulation. As shown in **Fig. 9**, these upregulated cytokines were GM-CSF, (common β chain family), IL-1α, IL-1β (IL-1 family), IL-6, LIF (IL-6/IL-12 family), IL-17A (IL-17 family), MIP1-α, MIP-1β, MIP-2, KC, LIX, MCP-1 (chemokines), G-CSF, VEGF (growth factors), and TNF-α (TNF family). In all these cases, the subpopulation with the highest cytokine concentration corresponded to mice with *F. animalis* gastric titers above 5×10^5^ CFU / g of stomach, and the cytokine concentration followed a positive linear correlation with the gastric *F. animalis* CFUs (**Supplementary Fig. 5**). For the other 15 quantified cytokines, which also included representatives of the IL-10, complement, interferon, and common γ chain families, we did not observe a significant difference between groups or a significant correlation with *F. animalis* CFUs (**Supplementary Table 1**). In summary, a high *F. animalis* infection load elicits the production of multiple proinflammatory cytokines and chemokines in metaplastic stomach tissues.

**Figure 9.**
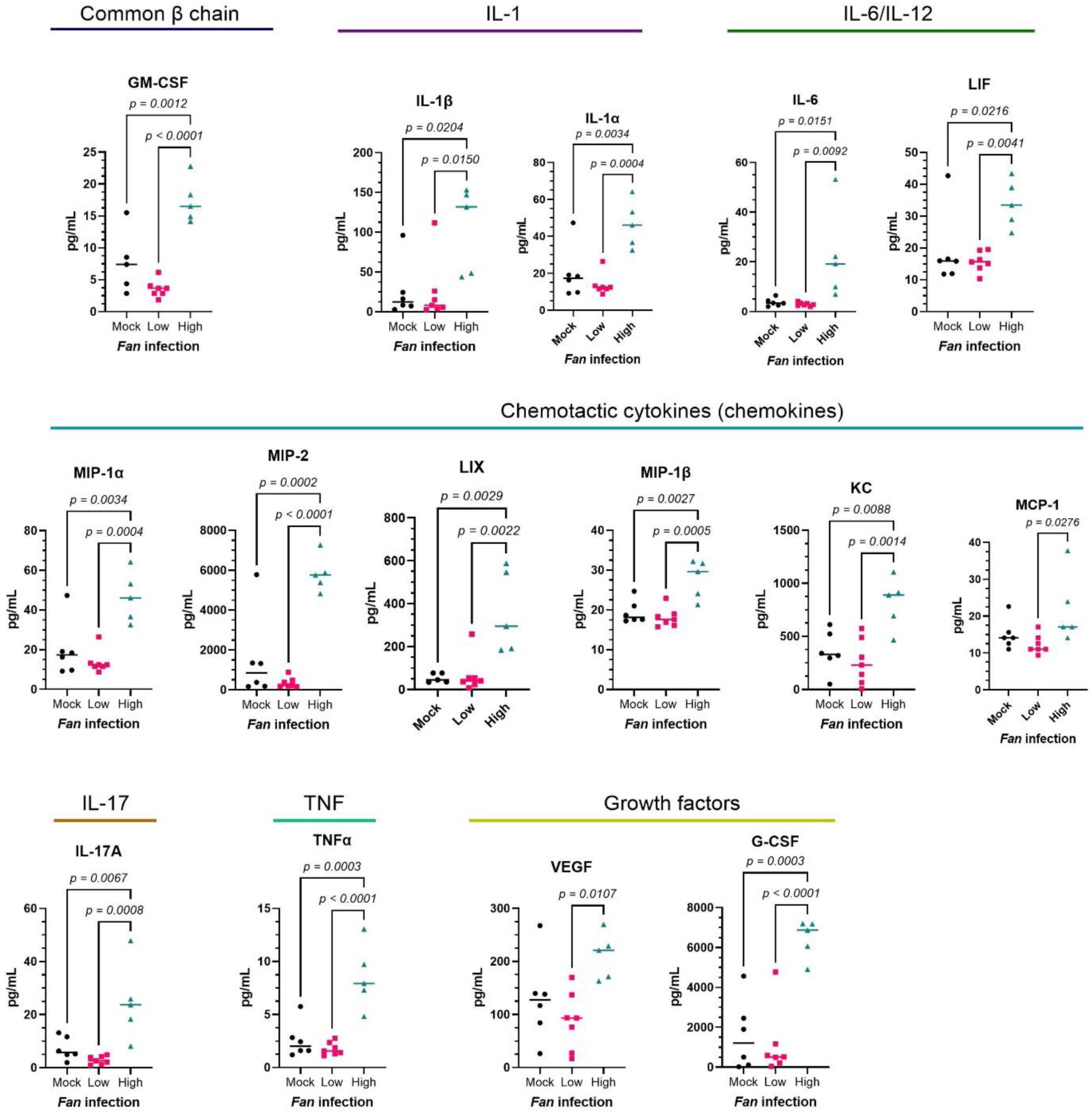
Concentration of proinflammatory cytokines measured in GIM stomach homogenates infected with *F. animalis*. *Mist1-Kras* mice were induced with tamoxifen and infected with *F. animalis* at the 6^th^ week post-induction. Stomachs were harvested and homogenized 6 weeks post-infection, and these homogenates were used for cytokine and CFU quantification. Data were classified into three groups according to their infection status: Mock (control), low, and high for mice with gastric *F. animalis* loads below and above 5×10^5^ CFU / g stomach, respectively. Horizontal lines represent the median value of each group. Data for each cytokine were analyzed through an ordinary one-way ANOVA to identify differences between group means, and the 15 cytokines shown in this figure correspond to those, out of 31, for which a statistically significant difference was found compared to the control group. Cytokine families are indicated above the plots. The *p*-values (only shown if < 0.05) were calculated from post-hoc Tukey’s multiple comparisons. This experiment was conducted in duplicate using 3 - 6 mice per group.

## DISCUSSION

Increasing evidence suggests that Fusobacteria possess cancer-promoting activities and colonize cancerous tumors, where they may exacerbate disease progression or influence response to treatment. We studied factors commonly driven by *H. pylori* chronic infection in the stomach that may enable Fusobacteria to colonize metaplastic gastric tissue and cancer cells. We found that both *F. animalis* and *F. nucleatum* adhere to and intracellularly invade human gastric adenocarcinoma cells (AGS) *in vitro*, with *F. animalis* exhibiting significantly higher survival, adherence, and invasion. Similarly, *F. animalis*, but not *F. nucleatum*, established stable gastric infections in the *Mist1-Kras* mouse model of GIM. Therefore, preneoplastic and cancerous gastric tissues are likely to select for species of Fusobacteria that carry virulence factors and other adaptations to the specific environmental stressors found in these environments, such as low pH, micronutrient availability, and immune response, among others. This is consistent with findings reporting that even though both species are facultative intracellular pathogens (46, 47), *F. animalis* dominates in colorectal cancer tumors (16). It is worth noticing that even though most literature on the Fusobacteria-cancer association focuses on *F. nucleatum*, molecular methods used when surveying patient samples (such as *nusG* or rDNA 16S amplification) often do not offer enough resolution to distinguish *F. nucleatum* from *F. animalis* or other species within the *sensu lato*.

The tumor microenvironment offers a low-oxygen niche for microbes. We found that hypoxia (1 % O_2_, 10 % CO_2_) supports more robust and prolonged infections by *F. animalis* in AGS cells, whereas microaerobic conditions (10 % O_2_, 10 % CO_2_) result in reduced survival and less stable infections. In this vein, Udayasuryan *et al.* 2024 (48) reported that hypoxia, in contrast to normoxia (18 % O_2_), favors infection by *F. nucleatum* in a colorectal cancer cell line (HTC116) and exacerbates the effects of low oxygen on the host’s gene expression. Altogether, these results suggest that hypoxia offers more biologically relevant conditions to test fusobacterial interactions with cancer cells, and thus should be prioritized in future research.

Infection by *H. pylori* often leads to disruption of the gastric microbiome (microecological dysbiosis), typically characterized by enrichment of oral bacteria, including *Fusobacterium* species, which may further persist as tumor-associated microbiota in cancer (11, 22, 43–45). Gastric acidity is a critical selection pressure for bacteria in the stomach (7, 49) that is altered by *H. pylori*-driven inflammation. Namely, reduction of the digestive enzyme- and acid-producing chief and parietal cells that occur during atrophic gastritis leads to elevated gastric pH (40). Zepeda-Rivera *et al.* 2024 reported that *F. animalis* exhibits higher pH stress tolerance compared to other *Fusobacterium* species, likely due to a putative glutamate-dependent acid resistance system (16). Similarly, Hara *et al.* 2024 observed an inverse correlation of Fusobacteria with the acidity of the tumor environment in esophagogastric junction and gastric cancer cases (20). According to our results in the mouse model, GIM onset enables *F. animalis* to colonize the gastric tissue, conditions under which an elevated pH facilitates, but does not determine infection. Thus, additional factors of the metaplastic gastric tissue are required for *F. animalis* growth.

We showed that *F. animalis* adherence and invasion of gastric cancer cells are primarily mediated by a Gal-GalNAc-binding mechanism, which is likely mediated by the autotransporter adhesin Fap2, a major virulence factor of the genus *Fusobacterium* (32) with dual activity of cell adhesion to Gal-GalNAc and of immune evasion via inactivation of NK cells (7). Lack of efficient techniques for genetic manipulation in fusobacterial species other than *F. nucleatum* still imposes a significant challenge to identifying the role of specific virulence factors. Recently, new methods have become available for *F. animalis*, which may help to overcome this limitation in future studies (50). Equally interesting, we observed a significant increase in gastric Gal-GalNAc expression in our mouse model from early stages of Kras-driven preneoplastic progression, but this did not entirely correlate with susceptibility to *F. animalis* infection, which did not occur until the 4^th^ week (*i.e.*, metaplasia onset). Hence, although Gal-GalNAc may promote fusobacterial infection, other metaplasia-related factors are still needed. Of the other factors we explored, neither metaplasia-driven inflammation nor concomitant *Hp* infection influenced *F. animalis* stomach colonization. Perhaps, the reduction in *H. pylori* abundance – reported as associated with an increase in Fusobacteria (11, 22, 43–45) – results from elimination by gastric inflammation controlling *H. pylori* infection over time (a phenomenon described by the hit-and-run model) (51) rather than bacteria-bacteria inhibitory interactions. Indeed, the predicted incidence of co-infection based on prevalence of each bacterium in their cohorts predicts at most a few co-infections. The lack of complete correlation between Gal-GalNAc production and fusobacterial infection in the murine GIM model and gastric cancer TMA suggests that while Gal-GalNAc may be a receptor that Fusobacteria exploit to invade cancer cells, it may not be the major factor promoting colonization of the tissue surface or mucus.

Unlike our findings in the AGS cell model, we did not find any instances of intracellular *F. animalis* in the mouse GIM model. Rather, we detected *F. animalis* growing in multispecies aggregates on the gastric mucus layer, which recapitulates the way Fusobacteria grow in the mouth, where they proliferate in complex multispecies biofilms on dental surfaces (52). In colorectal cancer, intratumoral *F. animalis* seems to be predominantly extracellular (53), and some studies have found that elevated pH may reprogram *F. nucleatum* gene expression to promote biofilm formation (54–56). Of note, we employed FISH as a method for localizing *F. animalis* in gastric tissue, which may technically limit detecting populations of intracellular bacteria. Relatedly, the Gal-GalNAc overproduction observed during gastric preneoplastic progression in the mouse model was primarily localized on the top 1/3 of the gastric glands, which comprises the pit and the isthmus regions, with the isthmus being the area with the major active stem cell population (57, 58). Hence, it is plausible that overexpression of Gal-GalNAc facilitates fusobacterial growth on the gastric mucosa and provides access to active host proliferative cells. Future research should seek to determine whether biofilms formed during gastric preneoplastic progression by fusobacteria and other secondary colonizers interact with gastric progenitor cell populations, as this could provide a mechanism to alter preneoplastic progression.

We found that the *F. animalis* – GIM tissue interaction elicited a pro-in ammatory response as measured by the production of 15 pro-inflammatory cytokines and chemokines in infections with high (above 10^5^ CFU/g) bacterial loads. Future research should validate the *F. animalis*-driven increase in the production of the specific cytokines identified in this screening. The type of upregulated cytokines and the cells producing them are important features of the pathogen-driven trajectory of disease progression.Our observations of elevated TNFα, interleukins (*i.e.,* IL1α/β and IL-6), and chemokines are consistent with those made in colorectal cancer cells and tissues (specifically to *F. animalis*) (17, 59), suggesting that the effect of *F. animalis* in gastric disease may underlie a similar mechanism of inflammation and immunosuppression as found in colorectal cancer. We did not identify the source of the upregulated cytokines, and since we did not detect instances of intracellular invasion in this model, future research should determine whether the interaction with the superficial mucosa is enough to trigger cytokine production by epithelial cells, or if a macrophage-mediated response is also involved, as reported in colorectal tumors (59, 60). Specifically, it has been reported that Fusobacteria may induce immunosuppression by recruiting myeloid cells via chemokine upregulation in macrophages while excluding and inhibiting T (59, 61) and NK cells (62).

Collectively, our results indicate that metaplasia onset renders the stomach susceptible to *F. animalis* infection. Even though metaplasia-induced Gal-GalNAc overproduction and reduced acidity facilitate growth, additional factors are required for infection. Altered mucin expression, gland atrophy, and the presence of other bacterial species are some variables not explored in this study that could contribute to fusobacterial gastric growth. While studies have identified Fusobacteria within gastric tumors, our work suggests that a tumor is not a necessary condition for colonization; indeed, *F. animalis* can colonize the metaplastic stomach environment and thus may contribute to tumor formation. Our results contrast with a prospective study of 44 gastric cancer cases that found that the presence of Fusobacteria was strongly correlated with poorer prognosis only in the diffuse type of gastric cancer, and the presence of Fusobacteria did not vary between gastritis and metaplastic lesions or tumors of the intestinal type of gastric cancer (23). Of note, gastric cancer is a highly heterogeneous disease driven by diverse genetic and environmental factors, leading to distinct phenotypes. AGS cells are derived from a case of intestinal-type gastric cancer, albeit lacking functional E-cadherin (63), a characteristic more commonly found in the diffuse type of gastric cancer. Likewise, the *Mist1-Kras* mouse model recapitulates the preneoplastic progression of the intestinal type of gastric cancer. Thus, future studies should investigate the interplay between diverse gastric cancer drivers and bacterial infections, including those established before neoplasia onset.

## MATERIALS AND METHODS

### Bacterial strains and culture conditions

*Fusobacterium animalis* SB010 (27) (donated by Dr. Susan Bullman) and *F. nucleatum* ATCC 23726 (donated by Dr. Chenggang Wu) were routinely grown in Fastidious Anaerobe Agar (Neogen) supplemented with 10 % v/v defibrinated horse blood (HemoStat) and JVN antibiotics (*i.e*., 3.0 μg/mL josamycin, 4.0 μg/mL vancomycin hydrochloride, and 1.0 μg/mL norfloxacin, all manufactured by Thermo Fisher) at 37 °C under anaerobic conditions using an AS-580 anaerobic chamber by Anaerobe Systems.

*Helicobacter pylori* PMSS1 (64) was cultured on Columbia agar plates (BD Biosciences) supplemented with 5 % v/v defibrinated horse blood (HemoStat) 0.2 % w/v β-cyclodextrin (Acros Organics), 10 μg/mL vancomycin (Sigma Aldrich), 5 μg/mL cefsulodin (Sigma Aldrich), 2.5 U/ml polymyxin B (Sigma Aldrich), 5 μg/mL trimethoprim (Sigma Aldrich), and 8 μg/mL amphotericin B (Sigma Aldrich). Plates were incubated at 37 °C under microaerobic conditions (10 % CO_2_ and 10 % O_2_) using a multi-gas incubator MCO-19M by Sanyo. Liquid cultures were grown in BB10: Brucella Broth supplemented with 10 % v/v fetal bovine serum (FBS, Corning).

### *Mist1-Kras* mouse model of gastric metaplasia

All mouse experiments described in this work were reviewed and approved by the Fred Hutchinson Cancer Center Institutional Animal Care and Use Committee (IACUC protocol No. 1531) and were performed in accordance with the recommendations in the National Institutes of Health (NIH) Guide for the Care and Use of Laboratory Animals. *Mist1-CreERT2^Tg/+^ LSL-Kras*^G12D*Tg/+*^ C57BL/6 mice (herein referred to as *Mist1-Kras* mice) have been previously described (36). Male and female mice aged 8 to 12 weeks were randomized to treatment groups. For *Kras*-driven metaplasia induction, mice received three subcutaneous doses of 5 mg of tamoxifen (Sigma Aldrich) in corn oil (Sigma Aldrich) with 10 % v/v ethanol over 3 days (delivered as a 50 µL injection per day). Control (sham-induced) mice were given a similar series of injections with corn oil only. All injected animals were monitored periodically until the injection lesions healed completely. Mice received gamma-irradiated rainbow foraging bits (Bio-Serv) after each injection and physical checkup as a means to soothe stress caused by the procedure. *Kras*+ mice correspond to mice at the 6-week post-induction timepoint unless otherwise stated.

### Infection of *Mist1-Kras* mice

For *Fusobacterium* infections, a single colony of the corresponding strain grown on JVN-supplemented agar was spread onto antibiotic-free FAA plates and incubated anaerobically for 2 days as described above. Cells were then resuspended and washed in phosphate-buffered saline (PBS). Mice were orally gavaged with 1 × 10^9^ colony-forming units (CFU) in 100 μL PBS. Mock-infected mice were gavaged with an equivalent amount of PBS.

For *H. pylori* infections, a similar protocol as that described above was followed with modifications: Mice were orally gavaged with 5 × 10^7^ mid-log liquid culture CFUs in 100 μL BB10, or mock-infected with an equivalent volume of BB10 only.

After the indicated infection period, mice were humanely euthanized by CO_2_ inhalation followed by cervical dislocation. Stomachs were aseptically harvested, and most of the non-glandular forestomach was resected (keeping the limiting ridge) and longitudinally cut in half. One section was homogenized in PBS and plated for CFU quantification. The remaining section was fixed in 10 % neutral-buffered formalin phosphate (Thermo Fisher) for two days. Fixed stomachs were then randomly grouped and embedded in paraffin blocks and cut into 4 μm sections on positively charged slides for further processing.

### Omeprazole treatment

Mice were orally gavaged daily with 40 μg of omeprazole per gram of body weight, prepared in 40 % PEG 400 in PBS. In an equivalent regime, control mice received 40 % PEG 400 in PBS (drug vehicle) only. Omeprazole and control treatments were initiated three days before infection with *F. animalis* and maintained throughout the duration of the experiment.

### Dexamethasone treatment

Dexamethasone sodium phosphate (Thermo Fisher) was added to drinking water at 1.2 mg/L and administered *ad libitum*. Dexamethasone administration was initiated at the 2^nd^ week after induction with tamoxifen and maintained until the end of the experiments. Water bottles containing dexamethasone were protected from light, and water was changed weekly. Since both changes in body weight and spleen weight are proxies for the dexamethasone effect, the body weight of the mice was monitored throughout the experiment. Spleens were harvested at the time of euthanasia to record their weight.

### Infection of AGS cells and confocal microscopy

The human gastric adenocarcinoma cell line AGS was acquired from ATCC and tested periodically for *Mycoplasma* by the Biospecimen Processing and Biorepository core at the Fred Hutchinson Cancer Center (Seattle, WA) with the MycoProbe Mycoplasma Detection Kit (R&D Systems). AGS cells were routinely cultured in DMEM10, which corresponds to DMEM (Gibco) supplemented with 10% v/v FBS (Corning) at 37 °C under microaerobic conditions of 10% CO_2_ in a 3110 Forma™ Series II CO_2_ incubator.

For infection assays, AGS cells were seeded at 4 × 10^5^ cells per well into 12-well plates and allowed to adhere overnight. 24 h cultures of *F. animalis* or *F. nucleatum* on FAA plates were resuspended and washed twice with PBS. Bacteria were diluted to the desired cell density in DMEM10 supplemented with 10 % BB10 and used to infect AGS cells. Unless otherwise stated, the corresponding *Fusobacterium* species was added at an MOI = 0.1 to the AGS cells and co-cultured under hypoxia (1.0 % O_2_, 10 % CO_2_) in a multi-gas incubator (MCO-19M – Sanyo). At the end of the co-culture incubation, supernatants were harvested to quantify free bacteria, wells were washed twice with PBS to remove loose bacteria, and total host-associated bacteria were quantified by lysing the AGS layers with ddH_2_O for 30 min and plating on FAA. To quantify intracellular bacteria, AGS cells were incubated with 300 μg/mL metronidazole (Sigma Aldrich) in DMEM10 with 10% BB10 for 1.5 h and washed twice with PBS before lysis.

For infection assays involving fluorescence imaging, cells were seeded onto 35 mm FluoroDish plates (WPI), and bacteria were added at MOI = 10. Bacteria were previously labeled with Carboxyfluorescein Succinimidyl Ester (CFSE, by Invitrogen) following the manufacturer’s instructions. After the co-culture period, infected cell layers were washed twice with PBS and fixed with 4 % paraformaldehyde in PBS for 10 min at room temperature. Cells were then washed with PBS twice, and permeabilized with blocking buffer (10 % wt/v BSA, 10 % v/v FBS, and 0.5 % v/v Triton X-100 in PBS) for 2 h at room temperature. After blocking, cells were washed twice with PBS and incubated with iFluor555-conjugated Phalloidin (AAT Bioquest) at 4 °C overnight. Finally, cells were washed with PBS twice and incubated with 1:1,000 DAPI (Invitrogen) at room temperature for 10 min before being mounted with ProLong Diamond Antifade Mountant (Invitrogen).

Samples were visualized with a Dragonfly 200 High-Speed Confocal Spinning disk confocal microscope using a 63X oil lens.

### GalNAc inhibition assay

*F. animalis cells* were preincubated anaerobically at 37 °C for 30 min with 25 mM GalNAc in the same medium used for infection before being used as described above. A control inoculum was preincubated in media without GalNAc. Both control and pretreated cells were plated to verify that GalNAc did not affect *F. animalis* viability. For imaging Gal-GalNAc production in non-infected AGS cells, the same protocol for fluorescence microscopy described above was followed using FITC-conjugated 50 µg/mL PNA (Invitrogen) during the first overnight incubation.

### *In situ* fluorescence hybridization (FISH)

This assay was carried out on paraffin-fixed gastric tissue sections of *F. animalis*- and mock-infected *Kras*+ mice using the RNAscope system (ACD-biotechne) in accordance with the manufacturer’s instructions with a Leica Bond RX autostainer. The RNAscope LS Multiplex Fluorescent Assay Kit was used for simultaneous assessment of *F. animalis* (rRNA 23S) and other Eubacteria (degenerate rRNA 16S) with the probes ACD 486418-C2 and ACD 464468-C3, respectively. In short, paraffin-embedded tissue sections were baked at 65 °C for 60 min, then deparaffinized and hydrated on the Leica Bond RX autostainer. Heat-Induced Epitope Retrieval was performed with reagents from the indicated RNAscope reagent kits according to the manufacturer’s instructions. Briefly, tissues were incubated in Tris/EDTA, pH 9.0, for 15 min at 95 °C, followed by a protein digestion in protease III for 15 min at room temperature. ISH probes and amplification reagents were applied following the manufacturer’s instructions. After staining, slides were manually dehydrated through graded alcohols, cleared in xylene, and mounted in Epredia Cytoseal XYL (Fisher).

### Gal-GalNAc quantification in mouse gastric tissue

Slides of paraffinized gastric tissue sections from mice at different points of the *Kras*-driven metaplastic cascade were deparaffinized using Histo-Clear solution (National Diagnostics) and rehydrated in sequential washes with decreasing concentrations of ethanol. The slides were then washed with PBS twice and permeabilized with blocking buffer (10 % wt/v BSA, 10 % v/v FBS, and 0.5 % v/v Triton X-100 in PBS) for 2 h at room temperature. After blocking, slides were washed twice with PBS and incubated with FITC-conjugated 50 µg/mL PNA (Invitrogen) at 4 °C overnight. The next day, slides were washed twice with PBS and incubated with 1:1,000 DAPI (Invitrogen) at room temperature for 10 min before being mounted with ProLong Diamond Antifade Mountant (Invitrogen). Slides were scanned with a SLIDEVIEW VS200 Universal Whole Slide Imaging Scanner (Evident) at 20X magnification.

To estimate Gal-GalNAc display on mouse stomach tissue, ImageJ 1.54 was used to measure the mean pixel intensity of the green channel (*i.e.*, PNA signal) of DAPI-positive area throughout the corpus tissue section of the scans described above. The corpus (region of interest) was delimited manually in each scan as the gastric tissue (between the submucosa and the mucus layer) between the first gland (limiting ridge) and the antrum (identified by the change in gland morphology to shorter gland height and different cell type composition).

### Tissue Microarray (TMA)

Generation and processing of the gastric cancer TMA utilized in this study are fully described in (38). In short, this TMA contains core sections representative of deep cancer, superficial cancer, and non-neoplastic tissue adjacent to cancer as classified by a pathologist. These samples were obtained from a cohort of 47 gastric cancer patients from medical centers in the U.S. Pacific Northwest. 1-2 unstained sections from each patient were used for DNA extraction, which was then tested by droplet digital PCR (ddPCR) to detect fusobacterial infection, amplifying a region of the *nusG* gene. The oligonucleotides used were: 5’-CAACCATTACTTTAACTCTACCATGTTCA-3’ (forward primer), 5’-GTTGACTTTACAGAAGGAGATTATGTAAAAATC-3’ (reverse primer), and GTTGACTTTACAGAAGGAGATTA (FAM probe). These probes have been previously used and validated for detecting Fusobacteria in clinical samples via qPCR (65). Samples with no detected copies were considered negative, and those with 1-2 copies / μL DNA were re-extracted and re-tested. A case was considered positive for fusobacterial infection if at least one core tested positive in the ddPCR assay.

Separately, TMA FFPE slides were stained with the FITC-conjugated PNA to detect Gal-GalNAc production following the protocol described above. Cases were classified as positive or negative for Gal-GalNAc display if at least one core showed FITC-PNA signal above the background level.

### Cytokine quantification

Stomach sections of *F. animalis*- and mock-infected *Kras*+ mice were homogenized in PBS, and total protein concentration was quantified by fluorometry in a Quibit machine (Thermo Fisher) and then diluted to the desired concentration in PBS supplemented with cOmplete EDTA-free protease inhibitor cocktail (Roche) and stored frozen. Cytokine quantification was carried out using the Mouse Cytokine/Chemokine 32-Plex Discovery Assay Array (MD32) by Eve Technologies Corporation (Canada). Samples were processed upon the first thaw.

### Statistical Analysis

Data plots and statistical tests were done using Prism 10.6.0. Effect size estimates were calculated manually using the *p*-values and descriptive statistics obtained with this software.

## ACKNOWLEDGMENTS

We thank Dr. Susan Bullman for providing *F. animalis* SB010, as well as for her critical advice, and all the support received from the Bullman lab with pilot experiments. We also thank Dr. Chenggang Wu for the kind donation of the *F. nucleatum* strains used in this study. We thank Jacob Frick, Armando Rodríguez, and Ciara Pike for their experimental assistance.

We thank Drs. Hoku West-Foyle, Julien Dubrulle, and Lena Schroeder in the Fred Hutch Cellular Imaging shared resource for their help with fluorescence microscopy assays and image analysis, and Paul Kong, from the Fred Hutch Experimental Histology shared resource, for his assistance with the mFISH assays.

This research was funded by the Gastric Cancer Foundation grant DMG 2021-024 and National Institutes of Health grants R01 AI54423 and R21 CA270512 (to N.R.S.) and R00 CA263036 (to VPO). This work was also supported by the Fred Hutchinson Cancer Center Microbiome Research Initiative Pilot Grant and the NIH P30 CA015704 of the Fred Hutch/University of Washington/Seattle Children’s Cancer Consortium, which includes the Cellular Imaging (RRID:SCR_022609) and Experimental Histopathology (RRID:SCR_022612) shared resources.

C. Gómez-Garzón is a Washington Research Foundation postdoctoral fellow.

## REFERENCES

1. Bray F, Laversanne M, Sung H, Ferlay J, Siegel RL, Soerjomataram I, Jemal A. 2024. Global cancer statistics 2022: GLOBOCAN estimates of incidence and mortality worldwide for 36 cancers in 185 countries. CA Cancer J Clin 74:229–263. 10.3322/caac.21708

2. de Martel C, Georges D, Bray F, Ferlay J, Clifford GM. 2020. Global burden of cancer attributable to infections in 2018: a worldwide incidence analysis. Lancet Glob Health 8:e180–e190. 10.1016/S2214-109X(19)30488-7

3. Correa P. 1988. A Human Model of Gastric Carcinogenesis. Cancer Res 48:3554–3560.

4. Duan Y, Xu Y, Dou Y, Xu D. 2025. *Helicobacter pylori* and gastric cancer: mechanisms and new perspectives. Journal of Hematology & Oncology 18:1–21. 10.1186/s13045-024-01654-2

5. Kusters JG, Van Vliet AHM, Kuipers EJ. 2006. Pathogenesis of *Helicobacter pylori* infection. Clin Microbiol Rev 19:449–490. 10.1128/cmr.00054-05

6. Suerbaum S, Michetti P. 2002. *Helicobacter pylori* Infection. New England Journal of Medicine 347:1175–1186. 10.1056/NEJMra020542

7. Beasley DE, Koltz AM, Lambert JE, Fierer N, Dunn RR. 2015. The Evolution of Stomach Acidity and Its Relevance to the Human Microbiome. PLoS One 10:e0134116. 10.1371/journal.pone.0134116

8. Gunathilake M, Lee J, Choi IJ, Kim Y Il, Kim J. 2021. Association between bacteria other than *Helicobacter pylori* and the risk of gastric cancer. Helicobacter 26:e12836. 10.1111/hel.12836

9. Jeong S, Liao YT, Tsai MH, Wang YK, Wu IC, Liu CJ, Wu MS, Chan TS, Chen MY, Hu PJ, Kao WY, Liu HC, Tsai MJ, Liu CY, Chang CC, Wu DC, Hsu YH. 2024. Microbiome signatures associated with clinical stages of gastric Cancer: whole metagenome shotgun sequencing study. BMC Microbiol 24:1–13. 10.1186/s12866-024-03219-2

10. Niikura R, Hayakawa Y, Nagata N, Miyoshi-Akiayama T, Miyabayashi K, Tsuboi M, Suzuki N, Hata M, Arai J, Kurokawa K, Abe S, Uekura C, Miyoshi K, Ihara S, Hirata Y, Yamada A, Fujiwara H, Ushiku T, Woods SL, Worthley DL, Hatakeyama M, Han YW, Wang TC, Kawai T, Fujishiro M. 2023. Non-*Helicobacter pylori* Gastric Microbiome Modulates Prooncogenic Responses and Is Associated With Gastric Cancer Risk. Gastro Hep Advances 2:684–700. 10.1016/j.gastha.2023.03.010

11. Hsieh YY, Tung SY, Pan HY, Yen CW, Xu HW, Lin YJ, Deng YF, Hsu WT, Wu CS, Li C. 2018. Increased Abundance of *Clostridium* and *Fusobacterium* in Gastric Microbiota of Patients with Gastric Cancer in Taiwan. Scientific Reports 8:1–11. 10.1038/s41598-017-18596-0

12. Urbaniak C, Cummins J, Brackstone M, Macklaim JM, Gloor GB, Baban CK, Scott L, O’Hanlon DM, Burton JP, Francis KP, Tangney M, Reida G. 2014. Microbiota of human breast tissue. Appl Environ Microbiol 80:3007–3014. 10.1128/aem.00242-14

13. Parhi L, Alon-Maimon T, Sol A, Nejman D, Shhadeh A, Fainsod-Levi T, Yajuk O, Isaacson B, Abed J, Maalouf N, Nissan A, Sandbank J, Yehuda-Shnaidman E, Ponath F, Vogel J, Mandelboim O, Granot Z, Straussman R, Bachrach G. 2020. Breast cancer colonization by *Fusobacterium nucleatum* accelerates tumor growth and metastatic progression. Nature Communications 2020 11:1–12. 10.1038/s41467-020-16967-2

14. Alkharaan H, Lu L, Gabarrini G, Halimi A, Ateeb Z, Sobkowiak MJ, Davanian H, Fernández Moro C, Jansson L, Del Chiaro M, Özenci V, Sällberg Chen M. 2020. Circulating and Salivary Antibodies to *Fusobacterium nucleatum* Are Associated With Cystic Pancreatic Neoplasm Malignancy. Front Immunol 11:552239. 10.3389/fimmu.2020.02003

15. Hayashi M, Ikenaga N, Nakata K, Luo H, Zhong PS, Date S, Oyama K, Higashijima N, Kubo A, Iwamoto C, Torata N, Abe T, Yamada Y, Ohuchida K, Oda Y, Nakamura M. 2023. Intratumor *Fusobacterium nucleatum* promotes the progression of pancreatic cancer via the CXCL1-CXCR2 axis. Cancer Sci 114:3666–3678. 10.1111/cas.15901

16. Zepeda-Rivera M, Minot SS, Bouzek H, Wu H, Blanco-Míguez A, Manghi P, Jones DS, LaCourse KD, Wu Y, McMahon EF, Park SN, Lim YK, Kempchinsky AG, Willis AD, Cotton SL, Yost SC, Sicinska E, Kook JK, Dewhirst FE, Segata N, Bullman S, Johnston CD. 2024. A distinct *Fusobacterium nucleatum* clade dominates the colorectal cancer niche. Nature 628:424–432. 10.1038/s41586-024-07182-w

17. Ye X, Wang R, Bhattacharya R, Boulbes DR, Fan F, Xia L, Adoni H, Ajami NJ, Wong MC, Smith DP, Petrosino JF, Venable S, Qiao W, Baladandayuthapani V, Maru D, Ellis LM. 2017. *Fusobacterium nucleatum* subspecies animalis influences proinflammatory cytokine expression and monocyte activation in human colorectal tumors. Cancer Prevention Research 10:398–409. 10.1158/1940-6207.capr-16-0178

18. Borozan I, Zaidi SH, Harrison TA, Phipps AI, Zheng J, Lee S, Trinh QM, Steinfelder RS, Adams J, Banbury BL, Berndt SI, Brezina S, Buchanan DD, Bullman S, Cao Y, Farris AB, Figueiredo JC, Giannakis M, Heisler LE, Hopper JL, Lin Y, Luo X, Nishihara R, Mardis ER, Papadopoulos N, Qu C, Reid EEG, Thibodeau SN, Harlid S, Um CY, Hsu L, Gsur A, Campbell PT, Gallinger S, Newcomb PA, Ogino S, Sun W, Hudson TJ, Ferretti V, Peters U. 2022. Molecular and Pathology Features of Colorectal Tumors and Patient Outcomes Are Associated with *Fusobacterium nucleatum* and Its Subspecies animalis. Cancer Epidemiology Biomarkers and Prevention 31:210–220. 10.1158/1055-9965.epi-21-0463

19. Liao Y, Wu YX, Tang M, Chen YW, Xie JR, Du Y, Wang TM, He YQ, Xue WQ, Zheng XH, Liu QY, Zheng MQ, Jia YJ, Tong XT, Zhou T, Li XZ, Yang DW, Diao H, Jia WH. 2024. Microbes translocation from oral cavity to nasopharyngeal carcinoma in patients. Nature Communications 2024 15:1 15:1–14. 10.1038/s41467-024-45518-2

20. Hara Y, Baba Y, Oda E, Harada K, Yamashita K, Toihata T, Kosumi K, Iwatsuki M, Miyamoto Y, Tsutsuki H, Gan Q, Waters RE, Komohara Y, Sawa T, Ajani JA, Baba H. 2024. Presence of *Fusobacterium nucleatum* in relation to patient survival and an acidic environment in oesophagogastric junction and gastric cancers. British Journal of Cancer 131:797–807. 10.1038/s41416-024-02753-0

21. Petkevicius V, Lehr K, Kupcinskas J, Link A. 2024. *Fusobacterium nucleatum*: Unraveling its potential role in gastric carcinogenesis. World J Gastroenterol 30:3972–3984. 10.3748/wjg.v30.i35.3972

22. Das A, Pereira V, Saxena S, Ghosh TS, Anbumani D, Bag S, Das B, Nair GB, Abraham P, Mande SS. 2017. Gastric microbiome of Indian patients with *Helicobacter pylori* infection, and their interaction networks. Scientific Reports 2017 7:1 7:1–9. 10.1038/s41598-017-15510-6

23. Boehm ET, Thon C, Kupcinskas J, Steponaitiene R, Skieceviciene J, Canbay A, Malfertheiner P, Link A. 2020. *Fusobacterium nucleatum* is associated with worse prognosis in Lauren’s diffuse type gastric cancer patients. Scientific Reports 2020 10:1 10:1–12. 10.1038/s41598-020-73448-8

24. Zepeda-Rivera MA, Dewhirst FE, Bullman S, Johnston CD. 2025. Addressing controversy in *Fusobacterium* nomenclature: what exactly does “*F. nucleatum*” refer to? Gut Microbes 17. 10.1080/19490976.2025.2514797

25. Chen S, Su T, Zhang Y, Lee A, He J, Ge Q, Wang L, Si J, Zhuo W, Wang L. 2020. *Fusobacterium nucleatum* promotes colorectal cancer metastasis by modulating *KRT7-AS*/KRT7. Gut Microbes 11:511–525. 10.1080/19490976.2019.1695494

26. Chen S, Zhang L, Li M, Zhang Y, Sun M, Wang L, Lin J, Cui Y, Chen Q, Jin C, Li X, Wang B, Chen H, Zhou T, Wang L, Hsu CH, Zhuo W. 2022. *Fusobacterium nucleatum* reduces METTL3-mediated m6A modification and contributes to colorectal cancer metastasis. Nature Communications 2022 13:1 13:1–16. 10.1038/s41467-022-28913-5

27. Bullman S, Pedamallu CS, Sicinska E, Clancy TE, Zhang X, Cai D, Neuberg D, Huang K, Guevara F, Nelson T, Chipashvili O, Hagan T, Walker M, Ramachandran A, Diosdado B, Serna G, Mulet N, Landolfi S, Ramon S, Fasani R, Aguirre AJ, Ng K, Élez E, Ogino S, Tabernero J, Fuchs CS, Hahn WC, Nuciforo P, Meyerson M. 2017. Analysis of *Fusobacterium* persistence and antibiotic response in colorectal cancer. Science 358:1443–1448. 10.1126/science.aal5240

28. Zhang T, Li Y, Zhai E, Zhao R, Qian Y, Huang Z, Liu Y, Zhao Z, Xu X, Liu J, Li Z, Liang Z, Wei R, Ye L, Ma J, Wu Q, Chen J, Cai S. 2025. Intratumoral *Fusobacterium nucleatum* Recruits Tumor-Associated Neutrophils to Promote Gastric Cancer Progression and Immune Evasion. Cancer Res 85:1819–1841. 10.1158/0008-5472.can-24-2580

29. Liu C, Yang Z, Tang X, Zhao F, He M, Liu C, Zhou D, Wang L, Gu B, Yuan Y, Chen X. 2023. Colonization of *Fusobacterium nucleatum* is an independent predictor of poor prognosis in gastric cancer patients with venous thromboembolism: a retrospective cohort study. Thromb J 21:1–11. 10.1186/s12959-022-00447-2

30. Hsieh YY, Tung SY, Pan HY, Chang TS, Wei KL, Chen WM, Deng YF, Lu CK, Lai YH, Wu CS, Li C. 2021. *Fusobacterium nucleatum* colonization is associated with decreased survival of *Helicobacter pylori*-positive gastric cancer patients. World J Gastroenterol 27:7311–7323. 10.3748/wjg.v27.i42.7311

31. Barranco SC, Townsend CM, Gasarteli C, Macik BG, Burger NL, Boerwinkle WR, Gourley WK. 1983. Establishment and Characterization of an in Vitro Model System for Human Adenocarcinoma of the Stomach. Cancer Res 43:1703–1709.

32. Abed J, Emgård JEM, Zamir G, Faroja M, Almogy G, Grenov A, Sol A, Naor R, Pikarsky E, Atlan KA, Mellul A, Chaushu S, Manson AL, Earl AM, Ou N, Brennan CA, Garrett WS, Bachrach G. 2016. Fap2 Mediates *Fusobacterium nucleatum* Colorectal Adenocarcinoma Enrichment by Binding to Tumor-Expressed Gal-GalNAc. Cell Host Microbe 20:215–225. 10.1016/j.chom.2016.07.006

33. Slade DJ. 2021. New Roles for *Fusobacterium nucleatum* in Cancer: Target the Bacteria, Host, or Both? Trends Cancer 7:185–187. 10.1016/j.trecan.2020.11.006

34. Brennan CA, Garrett WS. 2018. *Fusobacterium nucleatum* — symbiont, opportunist and oncobacterium. Nature Reviews Microbiology 2018 17:3 17:156–166. 10.1038/s41579-018-0129-6

35. Gibbs RJ, Chambers AC, Hill DJ. 2024. The emerging role of *Fusobacteria* in carcinogenesis. Eur J Clin Invest 54:e14353. 10.1111/eci.14353

36. Choi E, Hendley AM, Bailey JM, Leach SD, Goldenring JR. 2016. Expression of Activated Ras in Gastric Chief Cells of Mice Leads to the Full Spectrum of Metaplastic Lineage Transitions. Gastroenterology 150:918–930.e13. 10.1053/j.gastro.2015.11.049

37. O’Brien VP, Koehne AL, Dubrulle J, Rodriguez AE, Leverich CK, Kong VP, Campbell JS, Pierce RH, Goldenring JR, Choi E, Salama NR. 2021. Sustained *Helicobacter pylori* infection accelerates gastric dysplasia in a mouse model. Life Sci Alliance 4. 10.26508/lsa.202000967

38. O’Brien VP, Kang Y, Shenoy MK, Finak G, Young WC, Dubrulle J, Koch L, Rodriguez Martinez AE, Williams J, Donato E, Batra SK, Yeung CCS, Grady WM, Koch MA, Gottardo R, Salama NR. 2023. Single-cell Profiling Uncovers a Muc4-Expressing Metaplastic Gastric Cell Type Sustained by *Helicobacter pylori*-driven Inflammation. Cancer ResCommun 3:1756–1769. 10.1158/2767-9764.crc-23-0142

39. Tong QY, Pang MJ, Hu XH, Huang XZ, Sun JX, Wang XY, Burclaff J, Mills JC, Wang ZN, Miao ZF. 2024. Gastric intestinal metaplasia: progress and remaining challenges. J Gastroenterol 59:285–301. 10.1007/s00535-023-02073-9

40. Jencks DS, Adam JD, Borum ML, Koh JM, Stephen S, Doman DB. 2018. Overview of Current Concepts in Gastric Intestinal Metaplasia and Gastric Cancer. Gastroenterol Hepatol (N Y) 14.

41. Zhang X, Li C, Cao W, Zhang Z. 2021. Alterations of Gastric Microbiota in Gastric Cancer and Precancerous Stages. Front Cell Infect Microbiol 11:559148. 10.3389/fcimb.2021.559148

42. Ferreira RM, Pereira-Marques J, Pinto-Ribeiro I, Costa JL, Carneiro F, MacHado JC, Figueiredo C. 2018. Gastric microbial community profiling reveals a dysbiotic cancer-associated microbiota. Gut 67:226–236. 10.1136/gutjnl-2017-314205

43. Kamali N, Talebi Bezmin Abadi A, Rahimi F, Forootan M. 2025. *Fusobacterium nucleatum* confirmed in gastric biopsies of patients without *Helicobacter pylori*. BMC Res Notes 18:1–6. 10.1186/s13104-025-07165-8

44. Nascimento Araujo C do, Amorim AT, Barbosa MS, Alexandre JCPL, Campos GB, Macedo CL, Marques LM, Timenetsky J. 2021. Evaluating the presence of *Mycoplasma hyorhinis, Fusobacterium nucleatum, and Helicobacter pylori* in biopsies of patients with gastric cancer. Infect Agent Cancer 16:1–15. 10.1186/s13027-021-00410-2

45. Castaño-Rodríguez N, Goh KL, Fock KM, Mitchell HM, Kaakoush NO. 2017. Dysbiosis of the microbiome in gastric carcinogenesis. Sci Rep 7:1–9. 10.1038/s41598-017-16289-2

46. McGuire AM, Cochrane K, Griggs AD, Haas BJ, Abeel T, Zeng Q, Nice JB, Macdonald H, Birren BW, Berger BW, Allen-Vercoe E, Earl AM. 2014. Evolution of invasion in a diverse set of *Fusobacterium* species. mBio 5:e01864. 10.1128/mbio.01864-14

47. Umaña A, Sanders BE, Yoo CC, Casasanta MA, Udayasuryan B, Verbridge SS, Slade DJ. 2019. Utilizing whole *fusobacterium* genomes to identify, correct, and characterize potential virulence protein families. J Bacteriol 201:e00273–19. 10.1128/jb.00273-19

48. Udayasuryan B, Zhou Z, Ahmad RN, Sobol P, Deng C, Nguyen TTD, Kodikalla S, Morrison R, Goswami I, Slade DJ, Verbridge SS, Lu C. 2024. *Fusobacterium nucleatum* infection modulates the transcriptome and epigenome of HCT116 colorectal cancer cells in an oxygen-dependent manner. Communications Biology 2024 7:1 7:1–15. 10.1038/s42003-024-06201-w

49. Fisher L, Fisher A. 2017. Acid-Suppressive Therapy and Risk of Infections: Pros and Cons. Clinical Drug Investigation 2017 37:7 37:587–624. 10.1007/s40261-017-0519-y

50. Ko D, Garrett WS. 2025. Conjugation-based genetic manipulation of *Fusobacterium animalis*. mBio 10.1128/mbio.01714-25.

51. Hatakeyama M. 2014. *Helicobacter pylori* CagA and Gastric Cancer: A Paradigm for Hit-and-Run Carcinogenesis. Cell Host Microbe 15:306–316. 10.1016/j.chom.2014.02.008

52. Welch JLM, Rossetti BJ, Rieken CW, Dewhirst FE, Borisy GG. 2016. Biogeography of a human oral microbiome at the micron scale. Proc Natl Acad Sci U S A 113:E791–E800. 10.1073/pnas.1522149113

53. Luis J, Niño G, Ponath F, Ajisafe VA, Wargo JA, Johnston CD, Bullman S, Becker CR, Kempchinsky AG, Zepeda-Rivera MA, Gomez JA, Wu H, Terrazas JG, Bouzek H, Cromwell E, Chanana P, Wong M, Damania A, White MG, You YN, Kopetz S, Ajami NJ. 2025. Tumor-infiltrating bacteria disrupt cancer epithelial cell interactions and induce cell-cycle arrest. Cancer Cell 44:1–21. 10.1016/j.ccell.2025.09.010

54. Zilm PS, Mira A, Bagley CJ, Rogers AH. 2010. Effect of alkaline growth pH on the expression of cell envelope proteins in *Fusobacterium nucleatum*. Microbiology 156:1783–1794. 10.1099/mic.0.035881-0

55. Zilm PS, Bagley CJ, Rogers AH, Milne IR, Gully NJ. 2007. The proteomic profile of *Fusobacterium nucleatum* is regulated by growth pH. Microbiology 153:148–159. 10.1099/mic.0.2006/001040-0

56. Chew J, Zilm PS, Fuss JM, Gully NJ. 2012. A proteomic investigation of *Fusobacterium nucleatum* alkaline-induced biofilms. BMC Microbiol 12:1–14. 10.1186/1471-2180-12-189

57. Goldenring JR, Mills JC. 2017. Isthmus Time Is Here: Runx1 Identifies Mucosal Stem Cells in the Gastric Corpus. Gastroenterology 152:16–19. 10.1053/j.gastro.2016.11.028

58. Han S, Fink J, Jörg DJ, Lee E, Yum MK, Chatzeli L, Merker SR, Josserand M, Trendafilova T, Andersson-Rolf A, Dabrowska C, Kim H, Naumann R, Lee JH, Sasaki N, Mort RL, Basak O, Clevers H, Stange DE, Philpott A, Kim JK, Simons BD, Koo BK. 2019. Defining the Identity and Dynamics of Adult Gastric Isthmus Stem Cells. Cell Stem Cell 25:342–356.e7. 10.1016/j.stem.2019.07.008

59. Galeano Niño JL, Wu H, LaCourse KD, Kempchinsky AG, Baryiames A, Barber B, Futran N, Houlton J, Sather C, Sicinska E, Taylor A, Minot SS, Johnston CD, Bullman S. 2022. Effect of the intratumoral microbiota on spatial and cellular heterogeneity in cancer. Nature 2022 611:7937 611:810–817. 10.1038/s41586-022-05435-0

60. Kostic AD, Chun E, Robertson L, Glickman JN, Gallini CA, Michaud M, Clancy TE, Chung DC, Lochhead P, Hold GL, El-Omar EM, Brenner D, Fuchs CS, Meyerson M, Garrett WS. 2013. *Fusobacterium nucleatum* Potentiates Intestinal Tumorigenesis and Modulates the Tumor-Immune Microenvironment. Cell Host Microbe 14:207–215. 10.1016/j.chom.2013.07.007

61. Mima K, Sukawa Y, Nishihara R, Qian ZR, Yamauchi M, Inamura K, Kim SA, Masuda A, Nowak JA, Nosho K, Kostic AD, Giannakis M, Watanabe H, Bullman S, Milner DA, Harris CC, Giovannucci E, Garraway LA, Freeman GJ, Dranoff G, Chan AT, Garrett WS, Huttenhower C, Fuchs CS, Ogino S. 2015. *Fusobacterium nucleatum* and T Cells in Colorectal Carcinoma. JAMA Oncol 1:653–661. 10.1001/jamaoncol.2015.1377

62. Gur C, Ibrahim Y, Isaacson B, Yamin R, Abed J, Gamliel M, Enk J, Bar-On Y, Stanietsky-Kaynan N, Coppenhagen-Glazer S, Shussman N, Almogy G, Cuapio A, Hofer E, Mevorach D, Tabib A, Ortenberg R, Markel G, Miklić K, Jonjic S, Brennan CA, Garrett WS, Bachrach G, Mandelboim O. 2015. Binding of the Fap2 Protein of *Fusobacterium nucleatum* to Human Inhibitory Receptor TIGIT Protects Tumors from Immune Cell Attack. Immunity 42:344–355. 10.1016/j.immuni.2015.01.010

63. Busche S, Kremmer E, Posern G. 2010. E-cadherin regulates MAL-SRF-mediated transcription in epithelial cells. J Cell Sci 123:2803–2809. 10.1242/jcs.061887

64. Arnold IC, Lee JY, Amieva MR, Roers A, Flavell RA, Sparwasser T, Müller A. 2011. Tolerance Rather Than Immunity Protects From *Helicobacter pylori*–Induced Gastric Preneoplasia. Gastroenterology 140:199–209.e8. 10.1053/j.gastro.2010.06.047

65. Castellarin M, Warren RL, Freeman JD, Dreolini L, Krzywinski M, Strauss J, Barnes R, Watson P, Allen-Vercoe E, Moore RA, Holt RA. 2012. *Fusobacterium nucleatum* infection is prevalent in human colorectal carcinoma. Genome Res 22:299–306. 10.1101/gr.126516.111

