## Supplemental materials for "Metaplasia Enables Stomach Colonization by *Fusobacterium animalis*"

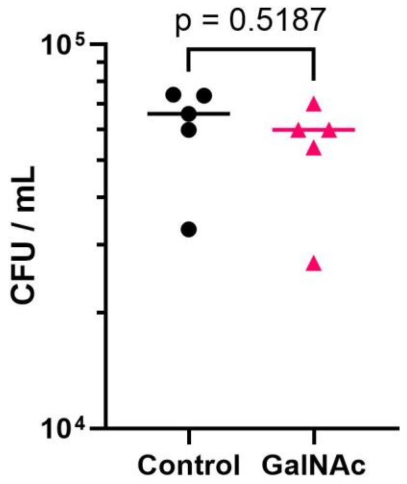


**Supplementary Figure 1. GalNAc does not affect the viability of *F. animalis*.** *F. animalis* CFU counts after 30 min incubation in 10:1 DMEM10 / BB10 medium supplemented with either 25 mM GalNAc or the equivalent amount of PBS (Control). These values correspond to independent experiments. Cell suspensions were prepared to achieve cell densities of ~$10^4$ CFU/mL, corresponding to an MOI of 0.1 in co-cultures with AGS cells. The *p*-value corresponds to an unpaired, two-tailed t-test between groups.

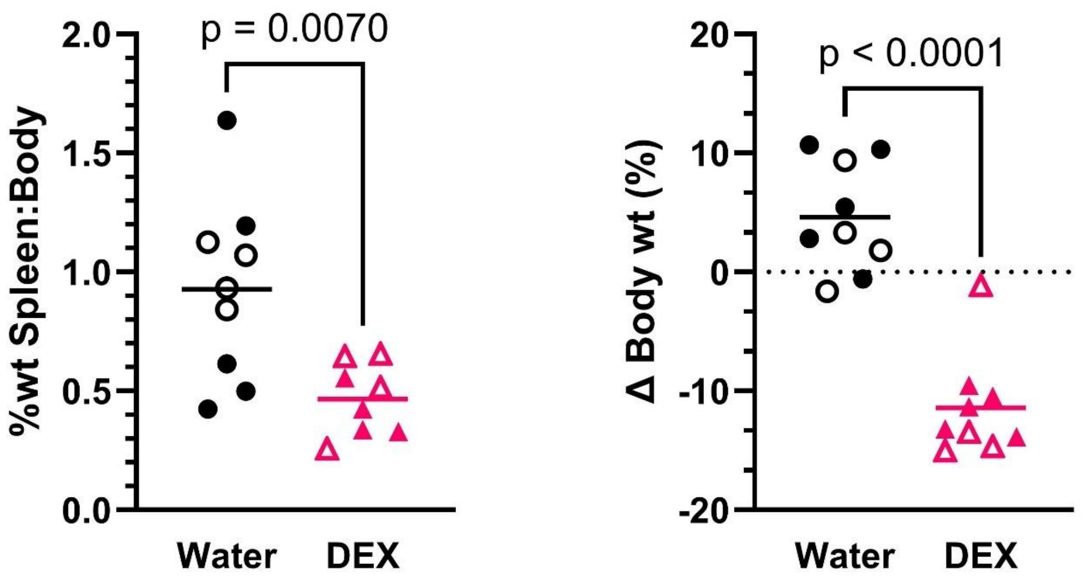


**Supplementary Figure 2. Body changes in mice treated with oral dexamethasone.** Comparisons were made between the control group that received non-acidified, autoclaved drinking water (Water) and the treatment group that received drinking water supplemented with 1 mg/L dexamethasone *ad libitum*. (DEX) **A)** Body weight (wt) percentage represented by the spleen at the end of the experiment. **B)** Percentage change in body weight between the start and the end of the experiment. Horizontal lines show the mean of each group, and *p*-values were calculated from non-paired, two-tailed t-tests. The data correspond to mice from the experiment shown in Fig. 4 of the main text.

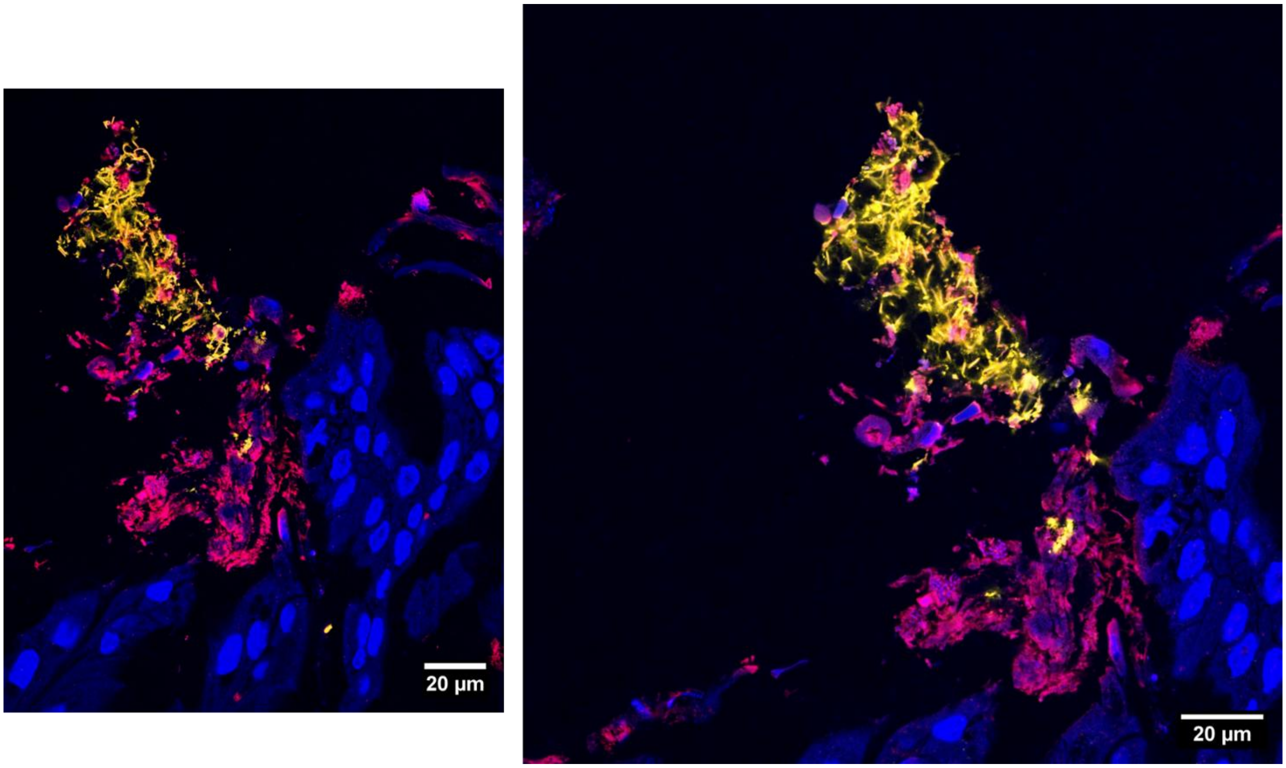


**Supplementary Figure 3. Magnification of a *Fan*-positive spot detected in a mFISH assay with a *Fusobacterium* 23S rRNA probe.** These images correspond to magnifications of the left panel of Fig. 8 in the main text. The foci labeled with the *Fusobacterium* probe (yellow) exhibits the classical spindle-like cell shape and size (2 – 10 µm) expected for *Fan*. Signal in red corresponds to Eubacteria 16S rRNA probe. Mouse tissue nuclei were labeled with DAPI (blue).

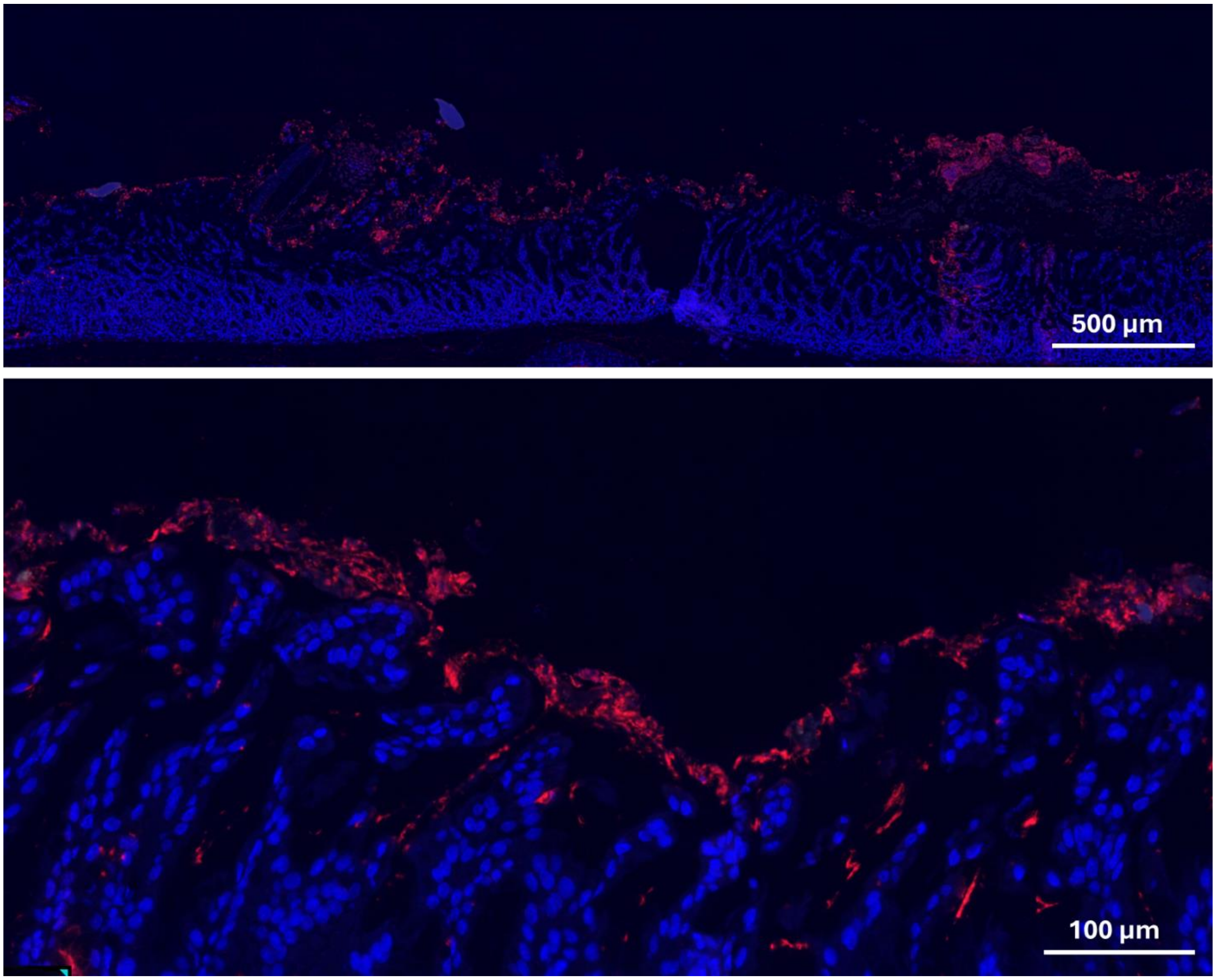


**Supplementary Figure 4. Control tissue for the mFISH assay shown described in Fig. 8 in the main text.** Both panels correspond to gastric tissue (corpus) of two representative mock-infected *Kras*+ mice labeled with the *Fusobacterium* 23S rRNA (yellow) and Eubacteria 16S rRNA (red) probes. Mouse tissue nuclei were labeled with DAPI (blue). No *Fan*-positive foci were detected in these samples, while similar patterns of Eubacteria-positive foci and gland atrophy were observed.

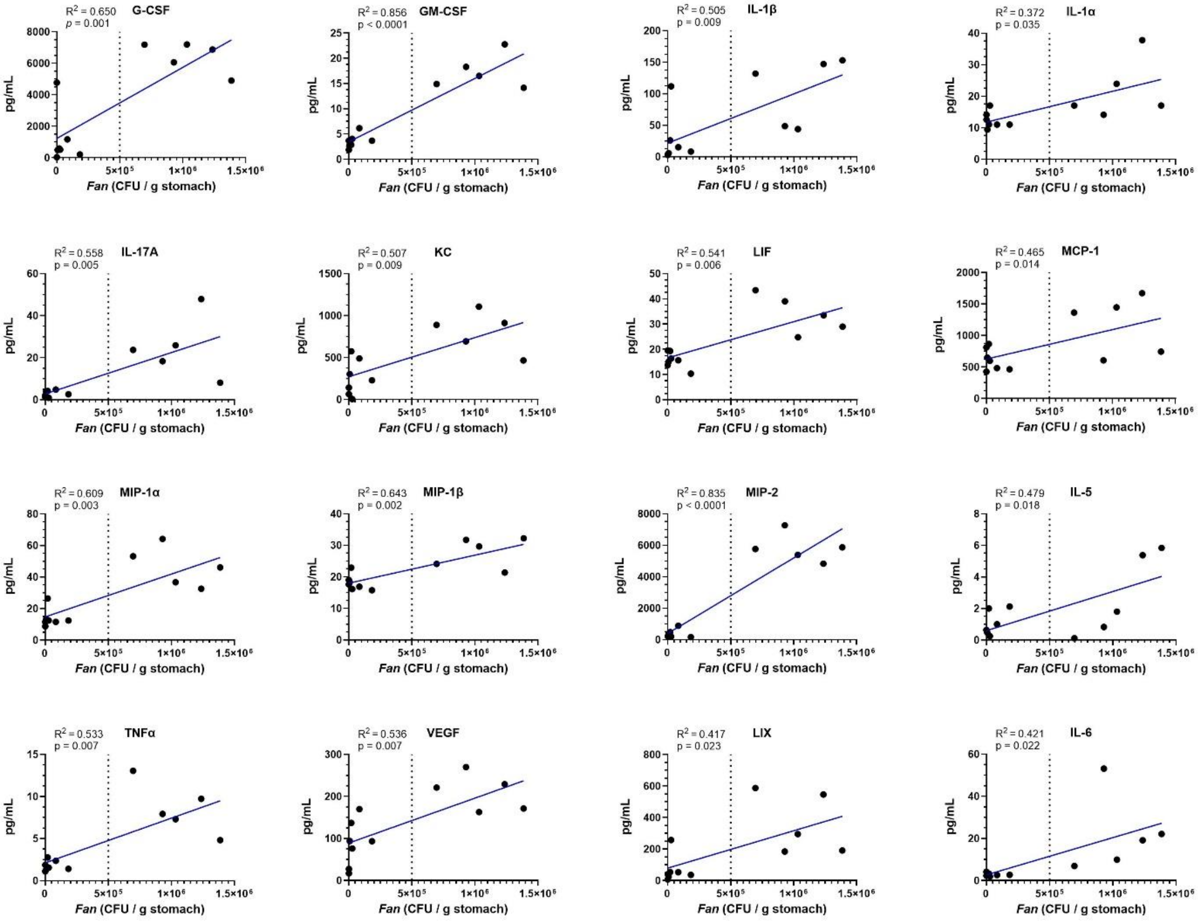


**Supplementary Figure 5. Linear regressions for cytokine concentration *vs. F. animalis* (*Fan*) titers for those cytokines with a significant difference between mock and infected groups.** The $R^2$ corresponds to the coefficient of determination, and the *p*-values were calculated from F tests for the slope equaling zero. The vertical dotted lines indicate the cutoff value ($5 \times 10^5$ CFU/g) used to define the low and high infection groups. Plotted data correspond to the experiment shown in Fig. 9 in the main text.

**Supplementary Table 1.** $R^2$ and *p*-values of the linear regressions of those quantified cytokines for which the null hypothesis of equal means could not be ruled out with α = 0.05 by an ANOVA test.

| Cytokine | *p*-value | $R^2$ |
|---|---|---|
| **Eotaxin*** | 0.022 | 0.424 |
| **IFNy** | 0.822 | 0.005 |
| **IL-2** | 0.904 | 0.001 |
| **IL-3** | 0.137 | 0.208 |
| **IL-4** | 0.087 | 0.265 |
| **IL-5**** | 0.018 | 0.479 |
| **IL-7** | 0.356 | 0.086 |
| **IL-9** | 0.303 | 0.105 |
| **IL-10** | 0.375 | 0.079 |
| **IL-12P40** | 0.397 | 0.072 |
| **IL-12P70** | 0.229 | 0.141 |
| **IL-15** | 0.656 | 0.021 |
| **IP-10** | 0.108 | 0.237 |
| **M-CSF** | 0.171 | 0.179 |
| **MIG** | 0.136 | 0.208 |
| **RANTES** | 0.918 | 0.001 |

* ANOVA *p*-value = 0.069, F = 3.19
** ANOVA *p*-value = 0.146, F = 2.22
